# Leucine zipper-based sorting system enables generation of multi-functional CAR T cells

**DOI:** 10.1101/2023.09.13.557232

**Authors:** Scott E. James, Sophia Chen, Brandon D. Ng, Jacob S. Fischman, Lorenz Jahn, Alexander P. Boardman, Adhithi Rajagopalan, Harold K. Elias, Alyssa Massa, Dylan Manuele, Katherine B. Nichols, Amina Lazrak, Nicole Lee, Teng Fei, Susan DeWolf, Jonathan U. Peled, Santosha A. Vardhana, Christopher A. Klebanoff, Marcel R. M. van den Brink

## Abstract

Resistance to chimeric antigen receptor (CAR) T cell therapy develops through multiple mechanisms including antigen-loss escape and tumor-induced immune suppression. Expression of multiple CARs may overcome multi-antigen-loss escape. Similarly, expression of switch receptors that convert inhibitory immune checkpoint signals into positive costimulatory signals may enhance CAR T cell activity in the tumor microenvironment. Engineering multiple features into one cell product, however, is limited by transgene packaging constraints of current vector systems. Here, we describe a leucine zipper-based cell sorting methodology that enables selective single-step immunomagnetic purification of cells co-transduced with two vectors, designed to potentially double the number of incorporated transgenes. This “Zip-sorting” system facilitated generation of T cells simultaneously expressing up to four CARs and co-expressing up to three switch receptors. These multi-CAR multi-Switch receptor arrays enabled T cells to eliminate antigenically heterogeneous syngeneic leukemia populations co-expressing multiple inhibitory ligands. Zip-sorted multi-CAR multi-Switch receptor T cells represent a potent therapeutic strategy to overcome multiple mechanisms of CAR T cell resistance.

---

Chimeric antigen receptor (CAR) T cell therapy has demonstrated remarkable therapeutic activity in refractory B cell malignancies and myeloma^1,2^. Despite this success, CAR T cell therapy faces a number of challenges that limit efficacy including CAR target antigen heterogeneity on tumor cells and associated antigen-loss escape, acquired CAR T cell dysfunction, and tumor-induced immune suppression^3–5^. These mechanisms of resistance to CAR T cell activity may occur concurrently, necessitating design of CAR T cells able to overcome multiple barriers to eliminate tumor cells.

Inter- and intra-patient heterogeneity of CAR target antigen expression represents a challenge in treating hematologic malignancies and solid tumors with CAR T cells, as entire populations or subsets of tumor cells may not express or may down-regulate selected target antigens^4,6–8^. Antigen-loss or antigen-low tumor escape occurs when target antigen expression on tumor cells is missing or insufficient to activate CAR T cell activity, respectively^4^. Dual-specificity CAR T cells have been evaluated as a strategy to address antigen heterogeneity and antigen-loss escape. However targeting more than two antigens may be required to eliminate malignancies with high antigen heterogeneity such as solid tumors^6,7^ and acute myeloid leukemia^8^. In preclinical models of B cell malignancies, engineered co-expression of CD19- and CD22-CARs prevented escape of leukemia with weak expression of both target antigens^9^. Similarly, CD123/CD19 dual-CAR and CD20/CD19 bi-specific tandem CAR T cells prevented escape of CD19-negative disease^10,11^. However, clinical trials have documented escape of antigen-low or multi-antigen-negative B cell leukemia and lymphoma populations following treatment with tandem CAR and dual-CAR T cells or sequential infusions of different single-antigen-targeting CAR T cells^12–15^. These findings suggest that T cells possessing greater than two antigen specificities may better resist escape of antigen-negative or antigen-low tumor cells.

The tumor microenvironment can impair CAR T cell activity via immune suppression mechanisms including upregulation of inhibitory immune checkpoint molecules on T cells and tumor cells and via promotion of T cell exhaustion^3,7,16,17^. In this setting, CAR T cell exhaustion develops in response to chronic antigen stimulation by tumor cells and is characterized by upregulation of inhibitory checkpoint molecules related to activation of transcription factor pathways including TOX^16–18^. CARs can also tonically signal in the absence of antigen stimulation, promoting inhibitory molecule expression and eventual exhaustion^19^. Strategies to overcome inhibitory receptor-associated immune suppression include co-expression of dominant negative or switch receptors that attenuate PD-1^20^, CD200R^21^, and Fas^22,23^ signaling or reprogram these inhibitory receptors to provide activating, rather than inhibitory costimulatory signals. Strategies to mitigate CAR T cell dysfunction resulting from tonic or antigen-triggered CAR signaling include temporal manipulation of CAR expression or activation^24^, optimization of CAR expression density^25^, attenuation of CAR CD3ζ signal strength^26^, and use of 4-1BB instead of CD28 costimulation^27^.

As cancer cells may utilize many immune evasion strategies simultaneously and CAR T cells may fail to eliminate malignancies due target antigen heterogeneity, T cell exhaustion, and tumor-induced immune suppression^3^, combinations of multiple synthetic biology solutions will likely be necessary to optimize CAR T cell activity. However, challenges emerge in attempts to combine multiple strategies, due to limitation in vector packaging capacity associated with larger vectors encoding multiple transgenes^28^. We present a leucine zipper-based cell sorting system that directs single-step immunomagnetic selection of cells transduced with two vectors with the goal of doubling the number of integrated transgenes. This system enables production of highly purified, complex cellular therapies utilizing clinically translatable magnetic sorting principles^29^. We utilized this system in syngeneic mouse models to engineer T cells co-expressing multiple CARs and switch receptors to overcome both antigen-loss escape and tumor-expressed inhibitory ligand immune evasion strategies.

## Results

### Leucine zipper-based sorting system enables single-step magnetic purification of dual-transduced cells

To generate multi-CAR T cells able to simultaneously target multiple antigens and overcome tumor-induced immune suppression strategies, we first developed a method to increase the number of transgenes integrated into engineered T cells by co-transducing with two vectors. Our goal was to isolate a uniform and highly purified T cell subpopulation transduced with two vectors using immunomagnetic selection, an approach compatible with clinical-grade cell production^29^. We constructed a sorting system (Fig. 1a-b) utilizing a heterodimerizing leucine zipper pair with each component encoded by one of two vectors: (Vector 1) a secreted affinity-tagged zipper able to bind an anti-tag magnetic bead for magnetic sorting and (Vector 2) a membrane-bound capture-zipper designed to retain the affinity-tagged zipper and magnetic bead uniquely on dual-transduced cells. Single-step immunomagnetic sorting would then selectively isolate dual-transduced cells (Fig. 1b).

**Fig. 1:**
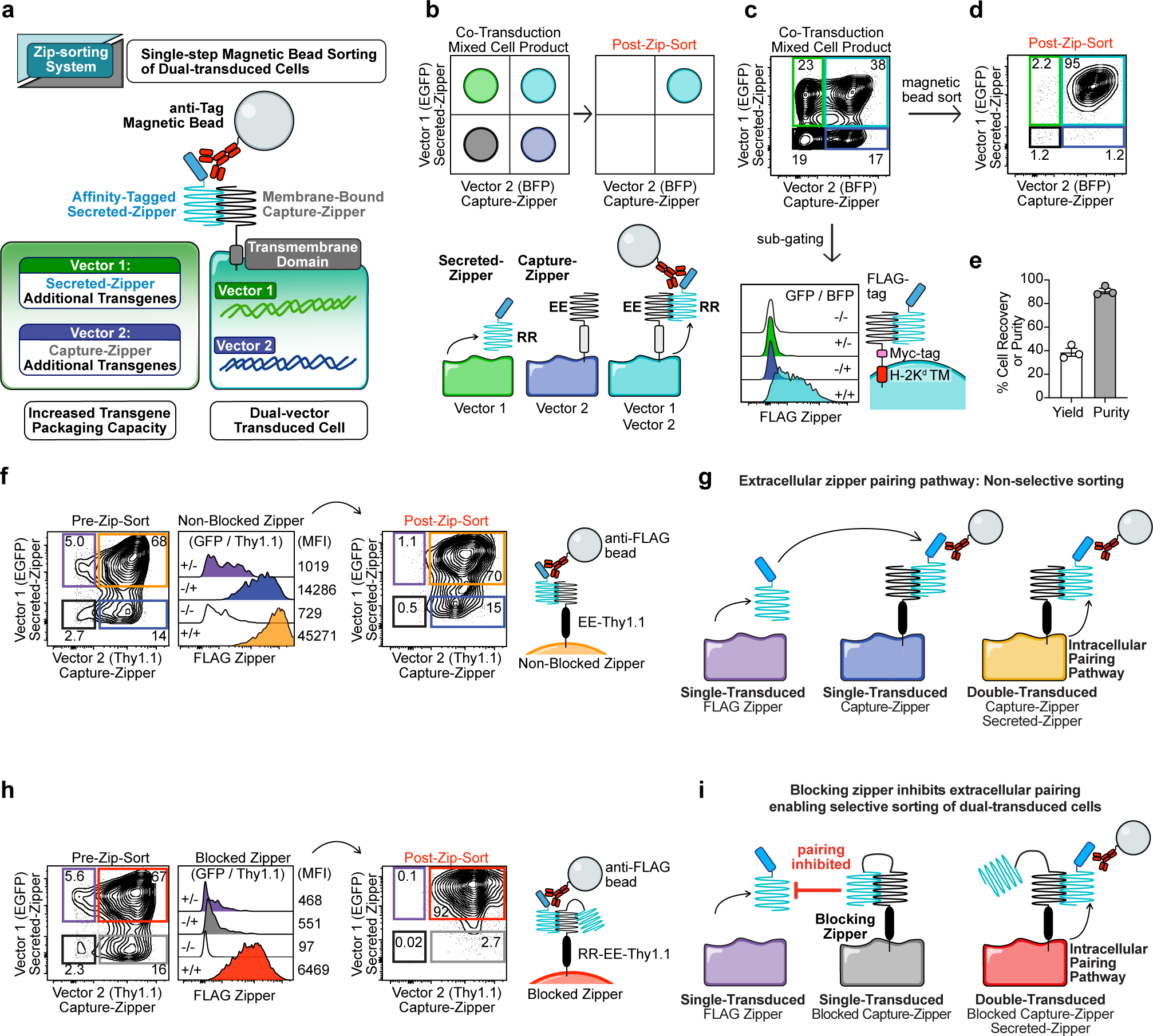
Engineering a leucine zipper-based cell sorting system. **a**, Leucine zipper sorting system principle. **b**, Predicted pre- and post-Zip-sorting results. **c**, Flow cytometry analysis of C1498 cells dual-transduced with FLAG-RR-CBR-GFP and EE-MHC-I-TM-P2A-BFP (see Extended Data Fig. 1 for vector maps). **d,** Zip-sort of purified cells from panel c. **e**, Yield and purity results from three Zip-sorts of cells described in panel c. Sort yield is defined as 100*(dual-transduced cells recovered/dual-transduced cells present pre-sort). Data are mean ±SEM (standard error of mean) of n=3 Zip-sorts. **f**, Flow cytometry analysis and Zip-sort results of C1498 cells co-transduced with FLAG-RR-CBR-GFP and EE-Thy1.1-P2A-BFP vectors. **g**, Proposed extracellular zipper pairing pathway. **h**, Flow cytometry analysis and Zip-sort results of C1498 cells co-transduced with FLAG-RR-CBR-GFP and RR-EE-Thy1.1-P2A-BFP vectors (performed at the same time as panel f). **i**, Proposed mechanism of blocking-zipper. Transduced cell lines were generated at least twice (independent transductions) with 1-2 Zip-sorts tested per replicate cell line.

To construct this system, we utilized a leucine zipper pair previously engineered for high-affinity heterodimerization^30^ via charge-based attraction of arginine (R) and glutamate (E) residues (abbreviated RR and EE, respectively; Supplementary Table 1). Prior to testing in primary T cells, we evaluated expression and function of this system by co-transducing C1498 leukemia cells with two vectors. Vector 1 encoded a secreted zipper (FLAG-RR) and an enhanced green fluorescence protein (EGFP) transduction reporter. Vector 2 encoded a membrane-tethered capture-zipper and a blue fluorescence protein (BFP) transduction reporter (Extended Data Fig. 1a). A previously described heterodimerizing leucine zipper CAR technology is based on extracellular pairing of exogenously supplied zipper-tagged scFvs with membrane bound zipper-based CARs^31^. We hypothesized that in contrast to the extracellular pairing observed in that system, co-expression of both an affinity-tagged-zipper and capture-zipper in the same cell might promote selective intracellular zipper pairing, thereby enabling preferential affinity-tag expression on dual-vector-transduced cells (Fig. 1b). As intended, FLAG-tagged zipper surface display occurred only on EGFP^+^ BFP^+^ C1498 cells dual-transduced with both zipper constructs (Fig. 1c). This selective pairing enabled single-step anti-FLAG bead immunomagnetic selection of dual-transduced cells with approximately 40% yield and 90% purity (Fig. 1d-e). We termed this methodology the Zip-sorting system and refer to zipper-based sorting as Zip-sorting.

### Blocking-zipper modification enables functional Zip-sorting system using transmembrane proteins with elongated extracellular domains

Fusing safety-switch molecules^32^, dominant negative receptors^20,22^, switch receptors^21,23^, costimulatory ligands^33^, or CARs with a Zip-sorting system capture-zipper component could improve the activity and safety of CAR T cells and enable purification of cells incorporating multiple transgenes. To investigate functionalizing these types of molecules with capture-zippers, we first evaluated construction of a safety-switch capture-zipper. On-target, off-tumor toxicity can impair the safety of CAR T cells and represents a challenge in targeting solid tumors^32^. Surface-expressed safety-switch constructs such as a truncated epidermal growth factor receptor (EGFRt)^34^ and a combined dual CD34/CD20 epitope tag stalk^35^ have been developed to enable immunomagnetic enrichment and antibody-mediated depletion of single-vector-transduced CAR T cells^32^. We constructed an analogous Zip-sorting safety-switch for use in syngeneic mouse models by appending the EE zipper to the extracellular domain of Thy1.1, which contains an epitope for anti-Thy1.1 antibody-mediated depletion (EE-Thy1.1, Extended Data Fig. 1a-b). In contrast to the original capture-zipper design lacking a spacer, EE-Thy1.1 strongly displayed the FLAG-RR zipper on both single- and dual-transduced EE-Thy1.1^+^ cells (Extended Data Fig. 1b), precluding selective sorting. This result suggested that capture-zippers with larger extracellular domains can pair extracellularly with secreted FLAG-RR zippers produced by other cells, similar to leucine zipper-based adapter CARs utilizing a CD8 hinge spacer^36^.

To enable selective sorting utilizing capture-zippers with larger extracellular domains, we hypothesized that further engineering of the EE-Thy1.1 zipper to include a linked RR blocking-zipper might inhibit extracellular pairing in single-transduced cells, while retaining intracellular pairing in dual-transduced cells (RR-EE-Thy1.1, Extended Data Fig. 1a, c-e and Fig. 1f-i). Indeed, single-transduced blocked zipper RR-EE-Thy1.1^+^ C1498 cells demonstrated an approximately 30-fold reduction in non-specific extracellular FLAG staining when co-cultured with FLAG-RR secreting C1498 cells (Extended Data Fig. 1c-d). However, dual-transduced FLAG-RR / RR-EE-Thy1.1^+^ C1498 cells demonstrated robust FLAG surface staining, which was approximately 10-fold higher than on single-transduced RR-EE-Thy1.1^+^ cells (Fig. 1h, red vs. gray histograms). This selective FLAG-zipper presentation enabled Zip-sorting of a highly purified EGFP^+^ Thy1.1^+^ dual-transduced population. In contrast, non-blocked EE-Thy1.1^+^ C1498 cells were non-selectively enriched for single and double-transduced cells (Fig. 1h vs. 1f, contour plots). Blocking-zipper mutants designed to repel the EE zipper via graded incorporation of repulsive glutamate and arginine substitutions demonstrated graded increases in extracellular zipper pairing (Supplementary Table 1 and Extended Data Fig. 1f; purple histograms and contour plots). These findings indicated that the blocking zipper inhibits extracellular pairing by competing with the FLAG-RR zipper for EE capture-zipper binding (Extended Data Fig. 1g).

Finally, to demonstrate that the blocking-zipper modification could promote selective sorting with other capture-zippers utilizing elongated extracellular domains, we evaluated blocked and non-blocked zippers appended to the human EGFRt safety-switch^34^ (Extended Data Fig. 1a). As with EE-Thy1.1, the non-blocked EE-EGFRt capture-zipper demonstrated extracellular pairing (Extended Data Fig. h, blue histogram). In contrast, the blocked RR-EE-EGFRt capture-zipper lacked detectable extracellular pairing (Extended Data Fig. 1h; gray vs. red histograms) and dual-transduced cells achieved sufficient FLAG surface expression for high purity Zip-sorting (Extended Data Fig. 1i). Together, these findings suggest that the blocking-zipper strategy may enable zipper functionalization of many types of synthetic biology constructs, facilitating purification of cells incorporating greater numbers of vector-encoded transgenes. In summary, we have developed a leucine zipper-based methodology to direct rapid and selective single-step isolation of dual-transduced cells using magnetic bead sorting, which is compatible with clinical applications^29^.

### Zip-sorted dual-CAR T cells prevent antigen-loss escape in vivo

To evaluate the Zip-sorting methodology in primary mouse T cells with the goal of preventing antigen-loss escape in a syngeneic dual-target antigen model, we generated a vector set encoding CD28-costimulated CD19- and CD20-targeting CARs (denoted CD1928z and CD2028z) and the Thy1.1 Zip-sorting / safety-switch components (Fig. 2a and Extended Data Fig. 2a). We additionally incorporated the inducible Caspase-9 safety-switch (iC9) for optional drug-mediated T cell depletion^37^, such that each vector encoded a safety-switch as an added safety feature (iC9 and RR-EE-Thy1.1; Extended Data Fig. 2a). Zip-sorting primary murine T cells achieved approximately 60% yield and over 90% purity of dual-vector-transduced T cells (Fig. 2b-c). Treatment with either the iC9 dimerizer AP20187 or with anti-Thy1.1 antibody plus complement efficiently depleted T cells, confirming activity of both safety-switch constructs (Fig. 2d-e). Dual-CAR T cells selectively lysed C1498 leukemia cells expressing cognate target antigens in vitro, demonstrating activity of both CARs (Fig. 2f).

**Fig. 2:**
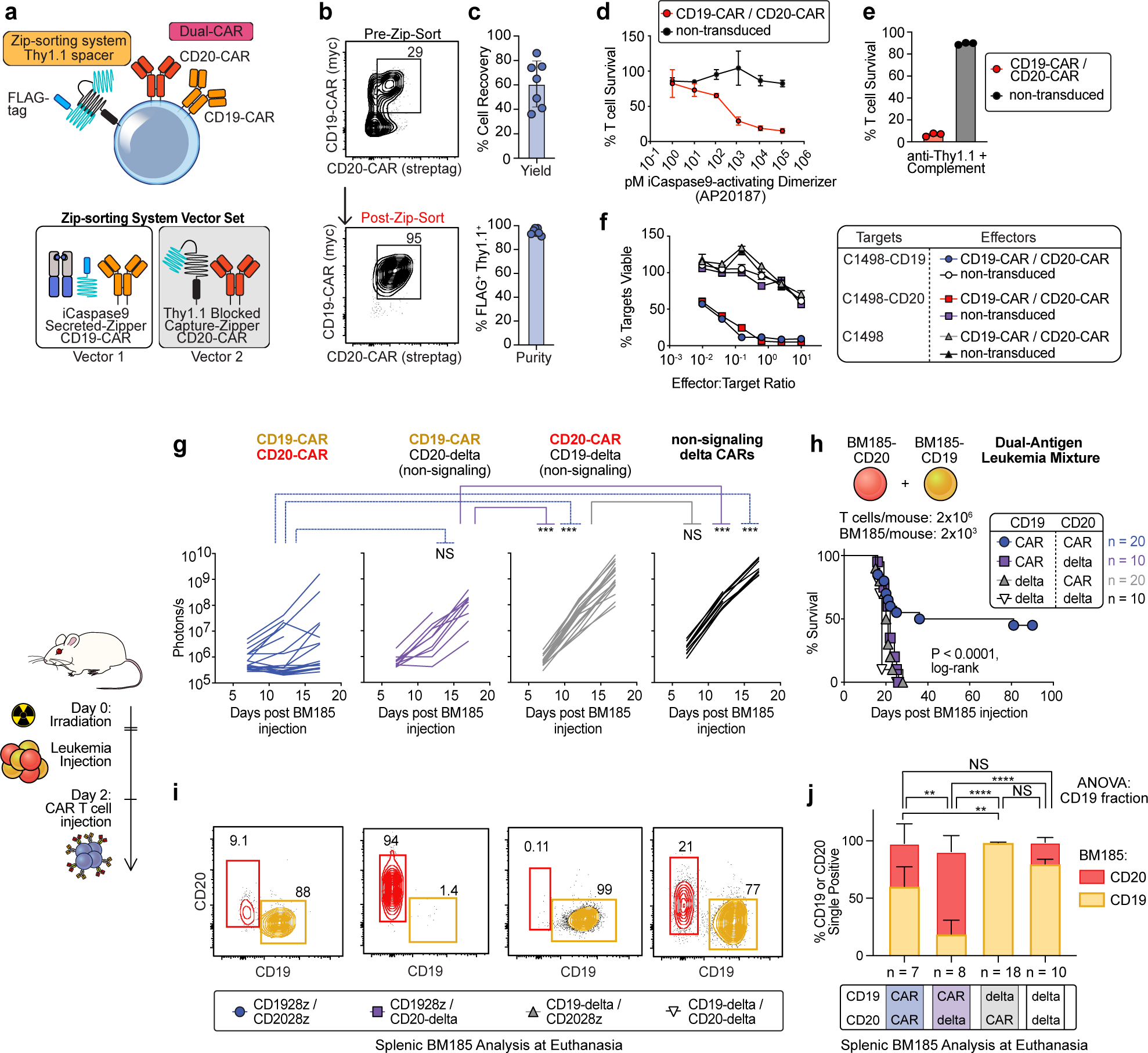
CD19/CD20 dual-CAR T cells prevent antigen loss escape in a heterogeneous leukemia model. **a**, Diagram of dual-CAR T cell design (see Extended Data Fig. 2a for vector maps). **b**, CAR expression of BALB/c primary T cells dual-transduced with FLAG-RR-iC9-CD1928z / RR-EE-Thy1.1-CD2028z vector set. **c**, Percent cell recovery (yield) and purity of Zip-sorted cells. n=7 biological replicates; mean ±SEM. **d**, Relative survival of Zip-sorted T cells treated with AP20187. Representative of n=2 donor experiments with triplicate wells; mean ±SEM. **e**, Relative survival of T cells incubated with complement and anti-Thy1.1 antibody for 30 minutes. Performed twice on T cells from one donor mouse with triplicate wells; mean ±SEM. **f**, 24h luciferase-based target lysis assay of Zip-sorted C57BL/6 (B6) dual-CAR T cells vs. C1498 targets. Representative experiment from n=2 donor experiments with triplicate wells; mean ±SEM. **g-j**, BM185-CD19/BM185-CD20 antigen-loss escape model (see Extended Data Fig. 2c for schematic). **g**, Leukemia BLI (ffluc) from four combined superimposed experiments. Log-transformed BLI values were compared using a Vardi test with false discovery rate (FDR) correction to compare area under the curve (AUC, see Methods). **h**, Survival, compared via log-rank test. **i**, Representative analysis of splenic BM185 populations at time of euthanasia for leukemia progression. **j**, Target antigen expression on splenic BM185 leukemia. % CD19^+^ fraction compared using one-way ANOVA with Tukey’s test for pairwise comparisons. *p≤0.05, **p≤0.01, ***p≤0.001, ****p≤0.0001, NS p>0.05.

To evaluate the potential of Zip-sorted dual-CAR T cells to eliminate a leukemia population with heterogeneous target antigen expression in vivo, we generated variants of the syngeneic BALB/c pre-B cell acute lymphoblastic leukemia cell line BM185 ffluc-Thy1.1-Neo^38,39^ modified to express the murine target antigens CD19 and/or CD20 (Extended Data Fig. 2b). We sublethally irradiated BALB/c mice as lymphodepleting preconditioning to enhance CAR T cell activity^40^. We then injected mice with a 1:1 mixture of BM185-CD19 and BM185-CD20 at a dose of 2×10^3^ cells/mouse and treated them 2 days later with 2×10^6^ Zip-sorted T cells expressing combinations of signaling-competent CD19 and CD20 CARs and signaling-incompetent CD3ζ-truncation delta CARs (Fig. 2g-j and Extended Data Fig. 2c). Zip-sorted dual-CAR T cells eliminated leukemia and produced long-term survival in a subset of mice, whereas CAR/delta, delta/CAR, and delta/delta T cells were unable to prevent progressive leukemia growth (Fig. 2g-h). Evaluation of splenic BM185 leukemia populations at the time of euthanasia for leukemia progression demonstrated selective outgrowth of the non-targeted BM185-CD19 or BM185-CD20 populations in recipients of single-antigen-targeting T cell recipients, consistent with antigen-loss escape (Fig. 2i-j). These findings demonstrated the ability of Zip-sorted dual-CAR T cells with dual-safety-switches to prevent antigen-loss escape in vivo.

### Culture with N-acetylcysteine or dasatinib enhances the anti-leukemia activity of dual-CAR T cells

To evaluate the capacity of Zip-sorted dual-CAR T cells to maintain activity against a higher leukemia burden, we evaluated dual-CAR T cells in mice injected with a 50-fold increased leukemia dose of 1×10^5^ BM185 cells/mouse (Extended Data Fig. 2d-f). In this setting, dual-CAR T cells produced modest anti-leukemia activity, yielding only a limited survival benefit (Extended Data Fig. 2d-f). CAR T cells can become exhausted in response to repetitive antigen encounter in the tumor microenvironment^16,17^. CAR T cells can also become dysfunctional as a result of tonic CAR signaling, which can chronically activate T cells in the absence of target antigen, resulting in loss of proliferative capacity and effector functions^19,24,25,27^. Additionally, pre-infusion CAR T cell products with increased exhaustion marker expression have been correlated with impaired clinical anti-leukemia activity^41^. We observed that resting, cultured Zip-sorted dual-CAR T cells upregulated the exhaustion-associated inhibitory receptor PD-1 in proportion to the number of CARs expressed (Extended Data Fig. 2g-h). The inhibitory receptor TIM-3 was upregulated similarly in CD8^+^ T cells expressing one or two CARs, while LAG-3 was upregulated most highly in CD4^+^ T cells with two CARs (Extended Data Fig. 2i), suggesting that dual-CAR tonic signaling promoted inhibitory receptor expression. Chronic T cell receptor signaling produces reactive oxygen species (ROS), which drive terminal T cell exhaustion and loss of proliferative capacity^42^. We also observed tonically increased total cellular ROS and mitochondrial superoxide in dual-CAR T cells (Extended Data Fig. 2j-m).

We hypothesized that scavenging ROS^42^ during T cell culture with N-acetylcysteine (NAC) or inhibiting tonic CAR signaling^24^ during culture with the Src kinase inhibitor dasatinib might prevent ROS upregulation and tonic signaling-induced dysfunction in dual-CAR T cells, thereby improving their activity in vivo. Therefore, we cultured dual-CAR T cells with 1 μM dasatinib or 10 mM NAC prior to infusion into mice. In contrast to control dual-CAR T cells, dasatinib- and NAC-cultured dual-CAR T cells eliminated an established 1×10^5^ BM185 leukemia cell dose in 100% of mice (Fig. 3a-b). Consistent with tonic CAR signaling, dual-CAR T cells demonstrated basal activation of NFAT, AP-1, and NFκB reporters (Fig. 3c). 24-hour culture with dasatinib inhibited tonic activation of these transcription factor pathways, while NAC mainly inhibited AP-1 and NFκB (Fig. 3c). The T cell exhaustion-associated hallmarks^43^ TOX and PD-1 were upregulated at baseline in dual-CAR T cells and increased further with target stimulation (Fig. 3d-e). However, culture with dasatinib or NAC inhibited tonic upregulation of multiple inhibitory receptors (Extended Data Fig. 3a-c). Additionally, culture with dasatinib inhibited exhaustion-associated modulation of TOX and TCF1 transcription factors^43^ in dual-CAR T cells (Fig. 3f). Dasatinib culture reduced total cellular ROS and mitochondrial superoxide in CD4^+^ and CD8^+^ dual-CAR T cells, while NAC only reduced mitochondrial superoxide in CD4^+^ T cells (Extended Data Fig. 3d-e), suggesting that dasatinib inhibits ROS production by blocking CAR signaling, while NAC enables T cells to tolerate elevated cellular ROS levels by increasing scavenging capacity^42^. Taken together, our findings suggest that in vitro culture with either NAC or dasatinib enabled our dual-CAR T cells to overcome development of ROS-associated tonic signaling-induced dysfunction during T cell culture and thereby eliminate larger leukemia populations in vivo.

**Fig. 3:**
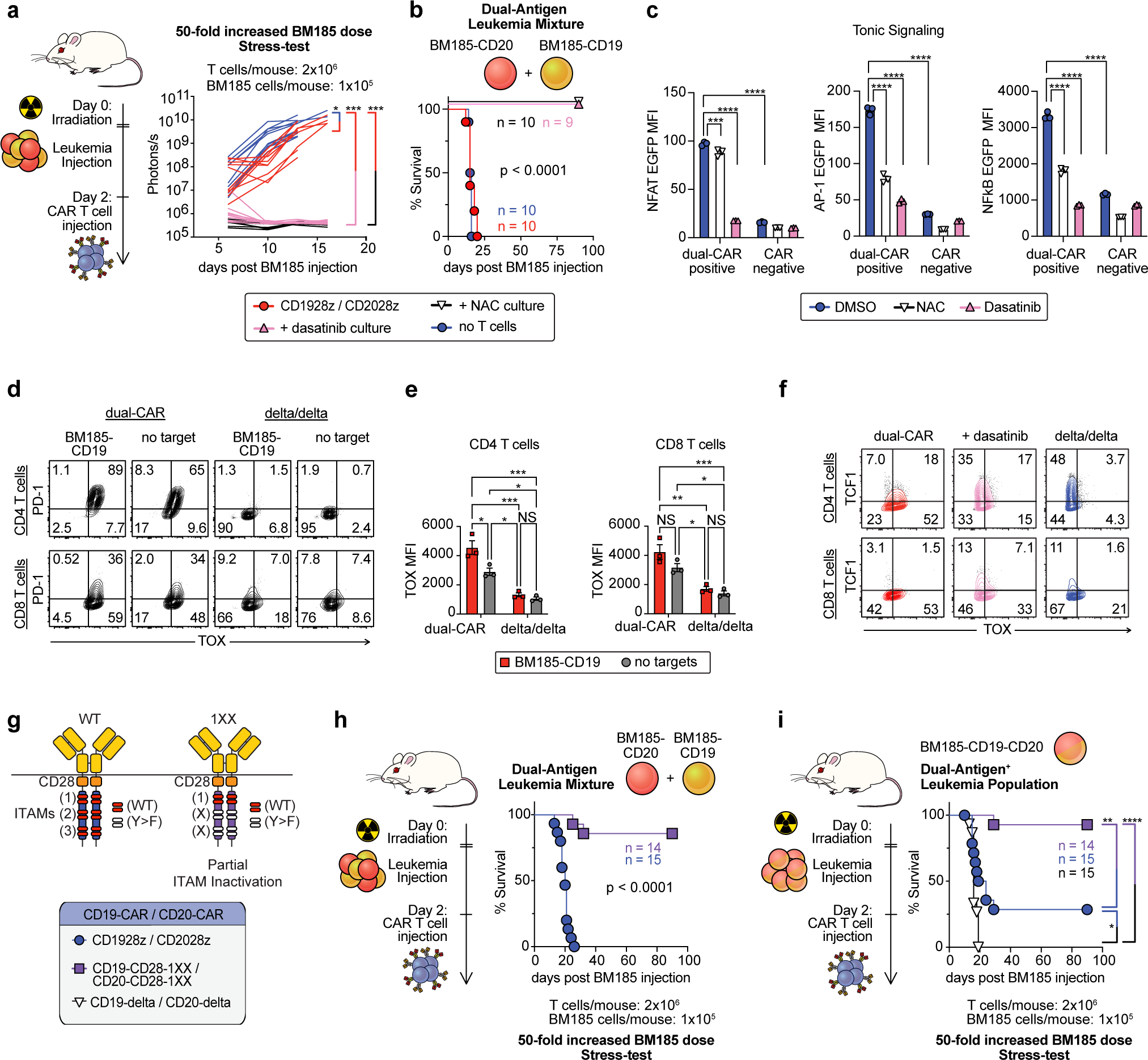
CAR T cell culture with NAC or dasatinib and ITAM attenuation enhances the anti-leukemia activity of dual-CAR T cells. **a-b.** BM185-CD19/BM185-CD20 antigen-loss escape model. BALB/c mice were treated with Zip-sorted dual-CAR BALB/c T cells cultured with 1 μM dasatinib (2 days), 10 mM NAC (3 days), or DMSO (3 days). **a**, Leukemia BLI from two combined experiments. Log-transformed BLI AUC values were compared using a Vardi test with FDR correction. **b**, Survival. **c**, NFAT, AP-1, or NFκB EGFP reporter analysis of unstimulated BALB/c T cells gated on dual-CAR or CAR-negative population following 24h culture with 1 μM dasatinib, 10 mM NAC, or DMSO. Representative of n=2 donor experiments, with mean ±SEM of triplicate wells. Statistical comparison via two-way ANOVA, with Tukey’s test for pairwise comparisons. **d-e**, PD-1 and TOX expression in Zip-sorted dual-CAR or delta/delta BALB/c T cells following 24h co-culture with BM185-CD19 or no targets. Mean ±SEM of triplicate samples were compared with a two-way ANOVA, with Tukey’s test for pairwise comparisons. **f**, Intracellular flow cytometry analysis of TCF1 and TOX expression in Zip-sorted dual-CAR or delta/delta BALB/c T cells cultured for 2 days with 1 μM dasatinib or DMSO. Representative experiment from n=2 donors**. g**, Diagram depicting ITAM mutations in 1XX CAR. **h**, Survival of BALB/c mice injected with BM185-CD19/BM185-CD20 (1:1) and treated with Zip-sorted dual-CAR WT CD3ζ or 1XX ITAM mutant dual-CAR T cells, from three combined experiments. **i**, Survival of BALB/c mice injected with dual-target-antigen expressing BM185-CD19-CD20 leukemia and treated with WT CD3ζ or 1XX dual-CAR T cells, from three combined experiments. Survival curves were compared via log-rank tests or pairwise log-rank test, with FDR correction. *p≤0.05, **p≤0.01, ***p≤0.001, ****p≤0.0001, NS p>0.05.

### CAR immunoreceptor tyrosine activation motif attenuation enhances the anti-leukemia activity of dual-CAR T cells

While culture of Zip-sorted dual-CAR T cells in dasatinib and NAC prevented the development of tonic signaling-induced dysfunction prior to infusion, T cell dysfunction can recur later in vivo in response to reestablishment of tonic signaling and in response to repetitive target antigen encounter^16,17,24^. Inactivation of selected CAR CD3ζ immunoreceptor tyrosine activation motifs (ITAMs) improves single-CAR T cell activity by attenuating CAR signal strength, thereby limiting terminal T cell differentiation and exhaustion^26^. We hypothesized that attenuation of CAR signaling via ITAM modulation might improve dual-CAR T cell activity against larger leukemia challenges. Dual-CAR T cells utilizing wild-type (WT) CD3ζ or the membrane-proximal single-ITAM CD3ζ variant 1XX (Fig. 3g) demonstrated equivalent CAR expression, target lysis in vitro, and inhibitory receptor expression (Extended Data Fig. 4a-c). However, 1XX ITAM mutant dual-CAR T cells exhibited reduced tonic production of total ROS and mitochondrial superoxide and maintained greater lytic activity following repetitive target stimulation in vitro (Extended Data Fig. 4d-f). 1XX dual-CAR T cells possessed greatly enhanced anti-leukemia activity compared with WT CD3ζ dual-CAR T cells, producing long-term survival in approximately 90% vs. 0% of mice in the higher-dose 1×10^5^ BM185 cells/mouse antigen-loss escape model (Fig. 3h). We additionally compared 1XX and WT dual-CAR T cells in recipients of BM185 leukemia co-expressing both CD19 and CD20 (BM185-CD19-CD20), designed to stress dual-CAR T cells further by simultaneously triggering both CARs. In this model, 1XX dual-CAR T cells retained enhanced activity, achieving long-term survival in nearly all mice (Fig. 3i). These findings indicate that ITAM attenuation improves the anti-leukemia activity of dual-CAR T cells.

### Co-expression of multiple switch receptors enhances the anti-leukemia activity of dual-CAR T cells

The tumor microenvironment can impair T cell function by inducing inhibitory molecule expression on T cells in response to repetitive antigen stimulation^16,17^ and by upregulating inhibitory ligands on tumor cells including FasL^22^, PD-L1^20^, and CD200^21^. We hypothesized that we could enhance the activity of multi-CAR T cells in vivo by antagonizing CAR-induced and tumor-expressed inhibitory checkpoint molecules (Fig. 4a) by co-expressing dominant negative receptor (DNR) or switch receptor (Fig. 4b) versions of the inhibitory receptors PD-1 and Fas. We also hypothesized that over-expressing the anti-apoptotic protein BCL-2 might enhance dual-CAR T cell survival and function. Each of these strategies has individually been reported to improve CAR T cell function and might achieve greater activity in combination^20,22,23,44^.

**Fig. 4:**
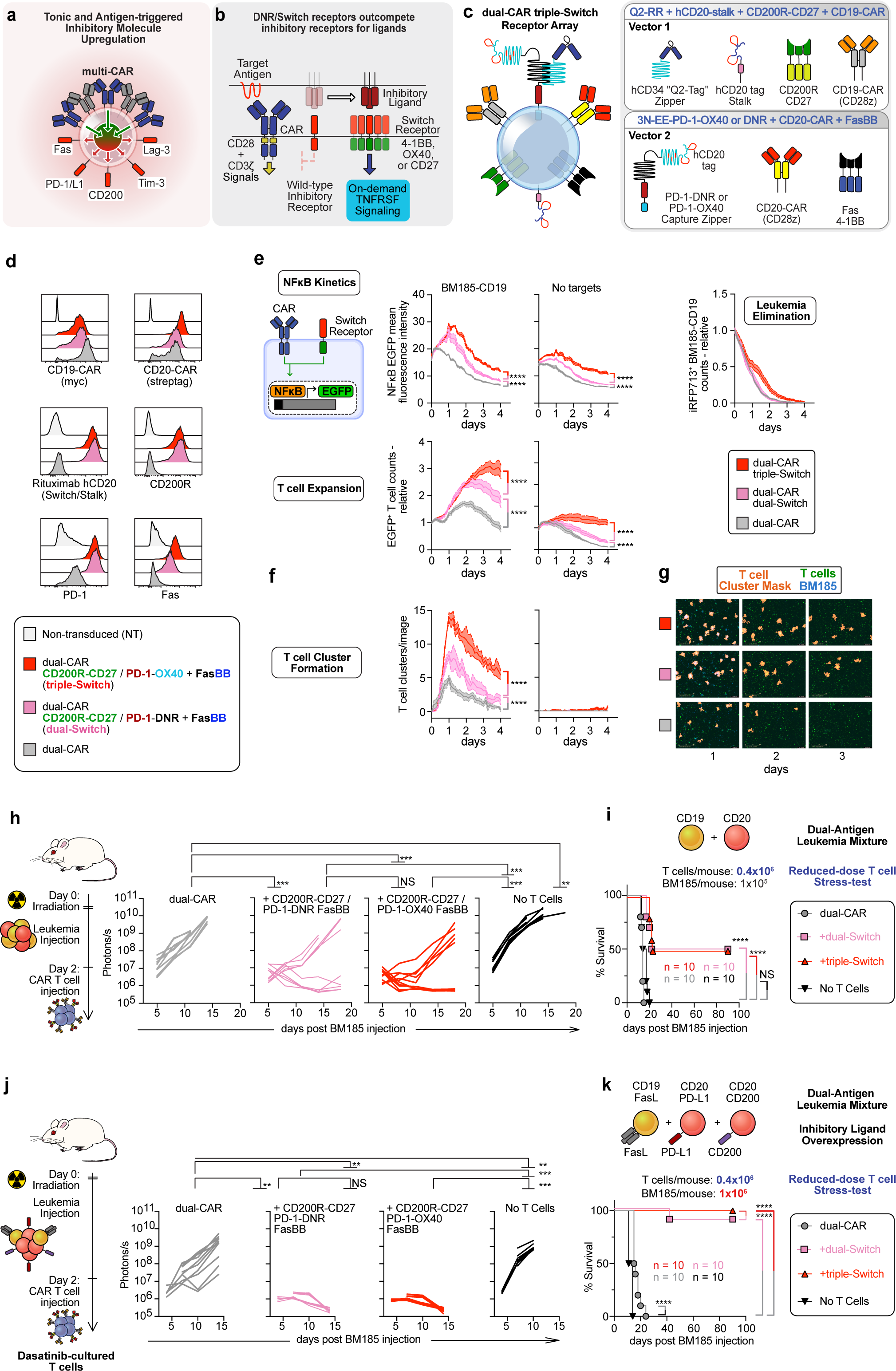
Co-expression of multiple switch receptors enhances the anti-leukemia activity of dual-CAR T cells. **a**, Diagram depicting CAR signaling-induced inhibitory molecule expression. **b**, Switch receptor principle. **c**, Diagram for dual-CAR multi-Switch receptor vectors (see Extended Data Fig. 6c for vector maps). R2-PD-1H-CD28TM-CD3ζΔ replaces iC9 as rituximab binding stalk for optional T cell depletion. **d**, Expression of CARs, safety-switches, and switch receptors in Zip-sorted BALB/c T cells. **e-g**. NFκB-reporter live-cell imaging (Incucyte) analysis of Zip-sorted BALB/c T cells stimulated with BM185-CD19-iRFP713 (E:T = 1:2) or unstimulated. **e**, (Top) Serial measurement of NFκB-EGFP reporter fluorescence intensity. (Bottom) EGFP^+^ T cell counts relative to initial values. (Right) iRFP713^+^ BM185-CD19 counts relative to initial values. **f**, Kinetic enumeration of large EGFP^+^ T cell clusters measuring at least 5000 μm^2^. Values represent mean ± SEM of triplicate samples, representative of n=3 donor experiments. AUC values were calculated in Prism and compared via one-way ANOVA, with Tukey’s test. **g**, Representative EGFP^+^ T cell cluster mask images of CAR T cells stimulated with BM185-CD19. **h-i**, BM185-CD19/BM185-CD20 antigen-loss escape model. Mice were treated with stress-test dose Zip-sorted dual-CAR BALB/c T cells. **h**, Leukemia BLI from two combined experiments. **i**, Survival. **j-k.** Sublethally irradiated BALB/c mice were injected with 8:1:1 mixture of BM185-CD19-FasL, BM185-CD20-PD-L1, and BM185-CD20-CD200 and treated with Zip-sorted dual-CAR BALB/c T cells, pre-cultured for 2 days in 1 μM dasatinib. **j**, Leukemia BLI from two combined experiments with different BLI imaging timing. **k**, Survival. BLI AUC were compared using a Vardi test with FDR correction. Survival differences were compared via pairwise log-rank test, with FDR correction. *p≤0.05, **p≤0.01, ***p≤0.001, ****p≤0.0001, NS p>0.05.

To encode multiple CARs, DNRs, and BCL-2 in two vectors, we designed a PD-1-DNR molecule incorporating a capture-zipper to enable Zip-sorting and simultaneously antagonize PD-L1/L2. We also included dual human CD20 (hCD20) mimotope tags^35^ as a safety-switch for optional rituximab antibody-mediated cell depletion (Extended Data Fig. 5a). This PD-1-DNR capture-zipper enabled high purity Zip-sorting using a CD34-tag^35^-based secreted-zipper, designed for use with CD34-microbeads utilized clinically to sort hematopoietic progenitor cells^45^ (termed Q2-RR, Extended Data Fig. 5a). We confirmed that the zipper-tagged PD-1-DNR maintained the ability to interact with PD-L1, by staining with soluble PD-L1 (PD-L1-Fc, Extended Data Fig. 5a), and by demonstrating that a zipper-tagged PD-1-CD28z receptor-based CAR promoted T cell lysis of BM185 cells overexpressing PD-L1 (Extended Data Fig. 5b-e). We incorporated the PD-1-DNR zipper, dominant negative Fas^22^ (FasDNR), and a caspase-cleavage resistant BCL-2^46^ into a CD1928z/CD2028z dual-CAR vector set and achieved high purity and expression of all constructs in Zip-sorted T cells (Extended Data Fig. 5a).

To determine if the dual-DNR + BCL-2 constructs could enhance the activity of dual-CAR T cells tonically upregulating inhibitory receptors, we utilized WT CD3ζ dual-CAR T cells cultured in the absence of NAC or dasatinib to treat mice injected with 1×10^5^ BM185 (Extended Data Fig. 5f-i). While dual-DNR-BCL2 dual-CAR T cells demonstrated transiently improved anti-leukemia activity, correlating with increased CD45.1^+^ congenic CAR T cell frequency in the blood, they did not improve overall survival of mice compared with dual-CAR T cells (Extended Data Fig. 5f-i). We therefore exchanged the FasDNR with a Fas-4-1BB switch receptor (FasBB)^23^, to provide supplemental 4-1BB costimulation. PD-1-DNR-BCL2-FasBB dual-CAR T cells promoted greater initial reduction in leukemia burden, higher day 10 CD45.1^+^ congenic CAR T cell frequency in the blood, and transiently improved mouse survival (Extended Data Fig. 5j-l). Still, we did not observe long-term survival, suggesting that these modifications were insufficient to maintain dual-CAR T cell activity against higher-dose BM185 leukemia challenge.

The improved survival of mice treated with FasBB^+^ dual-CAR T cells suggested that provision of additional costimulation via tumor necrosis factor receptor superfamily^47^ (TNFRSF) members such as 4-1BB might improve T cell activity and persistence. 4-1BB costimulation can promote long-term T cell survival^9,48^, improved T cell metabolism^49^, and resistance to development of T cell exhaustion^27^. We hypothesized that re-design of PD-1^50^ and CD200R1^21^ switch receptors to deliver “on-demand” TNFRSF signaling^33^ in response to upregulation of PD-L1 and CD200 inhibitory ligands expressed on T cells or tumor cells might enhance T cell expansion and survival. PD-L1 is upregulated on antigen-stimulated T cells, which impairs immune responses^51^. We also observed that the switch receptor FasBB strongly promoted upregulation of CD200 in antigen-stimulated dual-CAR T cells (Extended Data Fig. 6a-b). We designed the switch receptors PD-1-OX40 and CD200R-CD27 containing the costimulatory domains of the TNFRSF members OX40 and CD27^47^ to pair with FasBB, given synergistic T cell expansion reported to occur with combined 4-1BB and OX40 costimulation^52^ and with combination of OX40 or CD27 costimulation with PD-L1 blockade^53^.

We integrated PD-1-DNR, PD-1-OX40, and CD200R-CD27 into multi-Switch receptor configurations with FasBB and CD28-costimulated CARs and observed high expression of all receptors following Zip-sorting (Fig. 4c-d and Extended Data Fig. 6c). Expression of PD-1-DNR, PD-1-OX40, and CD200R-CD27 inhibited antigen-stimulated upregulation of PD-L1 and CD200 on dual-CAR T cells costimulated via FasBB, suggesting cognate receptor:ligand interactions^33,54^ (Extended Data Fig. 6d-g). Continuous live-cell microscopy imaging revealed that compared to dual-CAR T cells, dual-CAR dual-Switch (CD200R-CD27, PD-1-DNR, FasBB) and to an even greater extent, dual-CAR triple-Switch (CD200R-CD27, PD-1-OX40, FasBB) T cells activated larger peak antigen-stimulated NFκB responses, associated with greater T cell expansion (Fig. 4e). We also observed increased antigen-stimulated T cell cluster formation with multi-Switch CAR T cells, which has been correlated with NFκB-induced upregulation of the adhesion molecule ICAM-1^55,56^ (Fig. 4f-g and Supplementary Movie 1). In a reduced-dose “stress-test”, where we treated mice with only 0.4×10^6^ CAR T cells, dual-CAR dual-Switch and dual-CAR triple-Switch T cells exhibited greater anti-leukemia activity than dual-CAR T cells in the 1×10^5^ BM185 cell dose antigen-loss escape model, resulting in a subset of mice with long-term survival (Fig. 4h-i). Overall, these findings suggested that co-expression of multiple switch receptors enhances the activity of CD28-costimulated dual-CAR T cells by promoting T cell proliferation associated with augmentation of NFκB signaling.

### Multi-Switch receptor arrays enable dual-CAR T cells to eliminate leukemia populations expressing multiple inhibitory ligands

Malignant cells can express FasL^22^, PD-L1^20^, and CD200^21^ as immune evasion strategies. To simultaneously model antigen-loss escape and inhibitory ligand-based immune evasion strategies, we generated BM185 leukemia cells heterogeneously expressing CD19, CD20, and combinations of FasL, PD-L1, and CD200 (Extended Data Fig. 6h). CD1928z/CD2028z dual-CAR T cells cultured in dasatinib to reverse development of dysfunction during cell culture initially exerted anti-leukemia activity in mice injected with a 1:1:1 mixture of 1×10^5^ BM185-CD19-FasL, BM185-CD20-PD-L1, and BM185-CD20-CD200 cells. However, we observed eventual outgrowth of PD-L1-enriched leukemia in bone marrow, suggesting that PD-L1 overexpression inhibits dual-CAR T cell activity in vivo (Extended Data Fig. 6i-k).

To assess the potential of multi-CAR multi-Switch T cells to eliminate leukemia populations with antigen heterogeneity and inhibitory-ligand overexpression under stringent conditions, we evaluated a stress-test dose of 0.4×10^6^ dual-CAR T cells in an antigen-loss escape/inhibitory ligand expression model. Based on the reduced percentage of BM185-CD19-FasL observed in the BM of untreated mice (Extended Data Fig. 6k), we adjusted the ratio of BM185-CD19-FasL, BM185-CD20-PD-L1, and BM185-CD20-CD200 to 8:1:1 and increased the total BM185 dose 10-fold to 1×10^6^ cells/mouse. Dasatinib-cultured dual-CAR T cells were unable to control leukemia progression in this setting (Fig. 4j-k). In contrast, dasatinib-cultured dual-CAR multi-Switch T cells rapidly cleared leukemia, resulting in survival of 90-100% of mice. In an in vitro stress-test repetitive stimulation assay, dual-CAR triple-Switch T cells proliferated and eliminated BM185-CD19-FasL, whereas dual-CAR T cells were killed in response to FasL (Extended Data Fig. 6l). In response to BM185-CD20-PD-L1 and BM185-CD20-CD200, dual-CAR triple-Switch T cells demonstrated enhanced T cell expansion. Together, these results suggest that multi-Switch receptor arrays can enhance expansion of CD28-costimulated multi-CAR T cells to overcome both antigen-loss escape and inhibitory ligand-based immune evasion strategies.

### Multi-Switch receptor arrays enhance the activity of triple-CAR T cells

Acute myeloid leukemia^8^ and solid tumors^6^ exhibit antigen heterogeneity, suggesting that multi-antigen-targeting approaches may be needed to address these diseases. In clinical trials, CAR T cells sequentially or simultaneously targeting two B cell leukemia/lymphoma-associated antigens selected for outgrowth of target antigen-negative lymphoma populations^12,14,15^, suggesting that targeting more than two antigens may better resist this immune evasion mechanism. BAFF-R^57^ and CD79b^58^ have been validated as potential CAR targets for B cell malignancies in preclinical models. To model antigen-negative leukemia escape from CAR T cells due to high target antigen heterogeneity, we utilized the B6 genetic background acute myeloid leukemia cell line C1498 (expressing a CBR-hCD8-puro vector encoding click beetle red luciferase, human CD8 reporter, and puromycin resistance). C1498 does not express CD19, CD20, CD79b, or BAFF-R endogenously, facilitating development of a model for high antigen heterogeneity. We generated C1498 clones singly expressing each of these target antigens (Extended Data Fig. 7a). We then injected albino B6 mice with mixtures of C1498 clones singly expressing CD19 and CD20, or singly expressing CD19, CD20, CD79b, and BAFF-R and treated them with dasatinib-cultured 1XX-mutant CD19/CD20-targeting dual-CAR T cells (Extended Data Fig. 7b-c). We utilized albino B6 mice as lack of pigmentation facilitates bioluminescence imaging (BLI; Extended Data Fig. 7b). Dual-CAR T cells eliminated the CD19/CD20 dual-antigen leukemia mixture in vivo, producing long-term mouse survival, but they promoted outgrowth of C1498-BAFF-R and C1498-CD79b in mice receiving the quadruple-antigen leukemia mixture, consistent with immunoediting and antigen-loss escape (Extended Data Fig. 7d-e).

To eliminate a leukemia population with higher antigen heterogeneity, we designed a triple-CAR array targeting B cell malignancy-associated antigens CD20, CD79b, and BAFF-R (Fig. 5a and Extended Data Fig. 7f). To enhance the anti-leukemia activity of triple-CAR T cells in vivo, we combined three of the strategies that we found improved dual-CAR T cell activity: 1XX CD3ζ-based CARs, dasatinib culture, and multiple switch receptor co-expression. Zip-sorted triple-CAR and triple-CAR dual-Switch T cells (PD-1-OX40 and FasBB) highly expressed all encoded receptors (Fig. 5a-b). Zip-sorted triple-CAR T cells demonstrated potent lytic activity against cognate antigen-expressing C1498 targets in vitro (Extended Data Fig. 7g). In sublethally irradiated albino B6 mice injected with 1×10^6^ cells/mouse of a triple-antigen C1498 mixture (1:1:1 CD20^+^, CD79b^+^, BAFF-R^+^), dasatinib-cultured triple-CAR and triple-CAR dual-Switch T cells demonstrated potent anti-leukemia activity, achieving long-term survival of mice (Fig. 5c-d). Triple-CAR dual-Switch T cells demonstrated greater T cell BLI signal compared with triple-CAR T cells, indicating greater T cell expansion (Fig. 5e), and in a stress-test 0.4×10^6^ T cells/mouse dose challenge, triple-CAR dual-Switch T cells produced greater anti-leukemia activity (Fig. 5f-g). These results demonstrated the capacity for Zip-sorted multi-CAR T cells to eliminate leukemia populations with heterogeneous antigen expression, which was further enhanced by pairing with a multi-Switch receptor array.

**Fig. 5:**
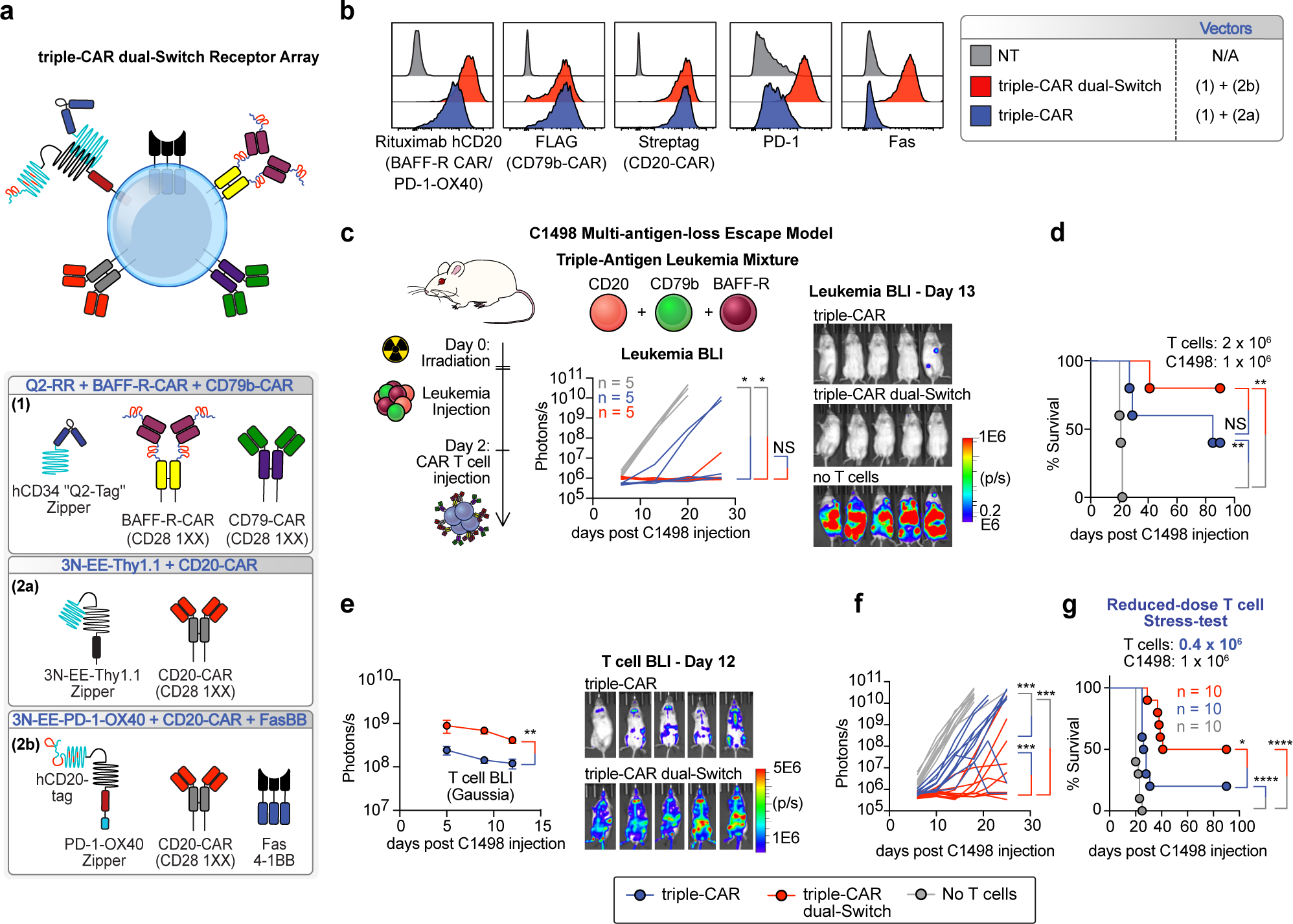
Multi-Switch receptor arrays enhance the activity of triple-CAR T cells. **a,** Diagrams for triple-CAR dual-Switch receptor array (see Extended Data Fig. 7f for vector maps). **b**, Flow cytometry analysis of Zip-sorted triple-CAR and triple-CAR dual-Switch albino B6 T cells. **c-e**. Sublethally irradiated albino B6 mice were injected with 1:1:1 ratio of C1498-CD20, C1498-CD79b, C1498-BAFF-R and treated with dasatinib-cultured triple-CAR and triple-CAR dual-Switch T cells co-transduced with a membrane-bound Gaussia luciferase (gLuc) vector: gLuc-PD-1H-CD24-GPI-P2A-EGFP for T cell BLI. **c**, C1498 BLI (CBR). **d**, Survival; from a single experiment. **e**, T cell BLI (Gaussia). **f-g**. Experimental setup as in panels c-e, but with stress-test 0.4×10^6^ T cell dose and T cell BLI not performed. **f**, C1498 BLI from two combined experiments. **g**, Survival. Log-transformed BLI AUC values were compared using a Vardi test with FDR correction. Survival differences were compared via pairwise log-rank test, with FDR correction. *p≤0.05, **p≤0.01, ***p≤0.001, ****p≤0.0001, NS p>0.05.

### Multi-Switch receptor arrays enhance the activity of quad-CAR T cells

To attempt to overcome even greater antigen heterogeneity and to demonstrate proof-of-principle of the increased transgene packaging capacity afforded by the Zip-sorting system, we next simultaneously targeted four tumor-associated antigens by co-expressing four CARs targeting CD19, CD20, CD79b, and BAFF-R (Extended Data Fig. 8a). To promote CD20-CAR expression in a larger vector also encoding a CD19-CAR and a capture-zipper, we constructed a smaller CD20-CAR using a single domain antibody (nanobody, abbreviated VHH, Extended Data Fig. 8a). However, the CD19-CAR expressed poorly from this vector (Extended Data Fig. 8a-b). As a result, quad-CAR T cells lysed C1498-CD19 targets weakly and promoted C1498-CD19 leukemia escape in vivo (Extended Data Fig. 8c-f). We therefore constructed a human CD19 (hCD19)-targeting CAR using a single domain antibody, as a mouse-CD19-targeting single domain antibody was not available, and paired it with the single domain CD20-CAR in a capture-zipper vector (Extended Data Fig. 9a). This Zip-sorted quad-CAR T cell configuration expressed all four CARs and lysed all cognate C1498 targets equally (Extended Data Fig. 9b-c). However, these quad-CAR T cells only briefly prolonged survival of mice bearing a quadruple-antigen mixture of C1498 targets (hCD19, CD20, CD79b, and BAFF-R; Extended Data Fig. 9d-e). T cell BLI on day 20 demonstrated a persistent CAR T cell bioluminescent signal and we did not observe selective outgrowth of C1498 leukemia antigen subsets (Extended Data Fig. 9f-g). These results argued against selective antigen loss escape, suggesting that these quad-CAR T cells were globally dysfunctional or insufficiently active

To overcome the poor activity of these quad-CAR T cells, we co-expressed multi-Switch receptor arrays. We designed a quad-CAR vector set combining four CARs and a CD200R-CD27 switch-receptor capture-zipper and co-transduced T cells with a third non-zipper vector encoding PD-1-OX40 and FasBB switch receptors (Fig. 6a and Extended Data Fig. 9a). Following Zip-sorting, all receptors expressed highly (Fig. 6b). In the setting of an established C1498 quadruple-antigen leukemia mixture in vivo, quad-CAR dual-Switch (PD-1-OX40 and FasBB) and triple-Switch (CD200R-CD27, PD-1-OX40, and FasBB) T cells produced greater overall T cell BLI signals in vivo, compared with quad-CAR T cells, and demonstrated enhanced BLI signals over the femurs, suggesting greater bone marrow T cell expansion (Fig. 6c-e). Multi-Switch quad-CAR T cells mediated superior anti-leukemia activity compared with quad-CAR T cells, achieving long-term survival in a subset of mice, and demonstrating the in vivo activity of CAR T cells able to simultaneously target four antigens (Fig. 6f-g). Together, these findings suggest that co-expression of multiple switch receptors promotes the proliferation and function of Zip-sorted multi-CAR T cells to enable elimination of leukemia populations with highly heterogeneous antigen expression (Extended Data Fig. 10).

**Fig. 6:**
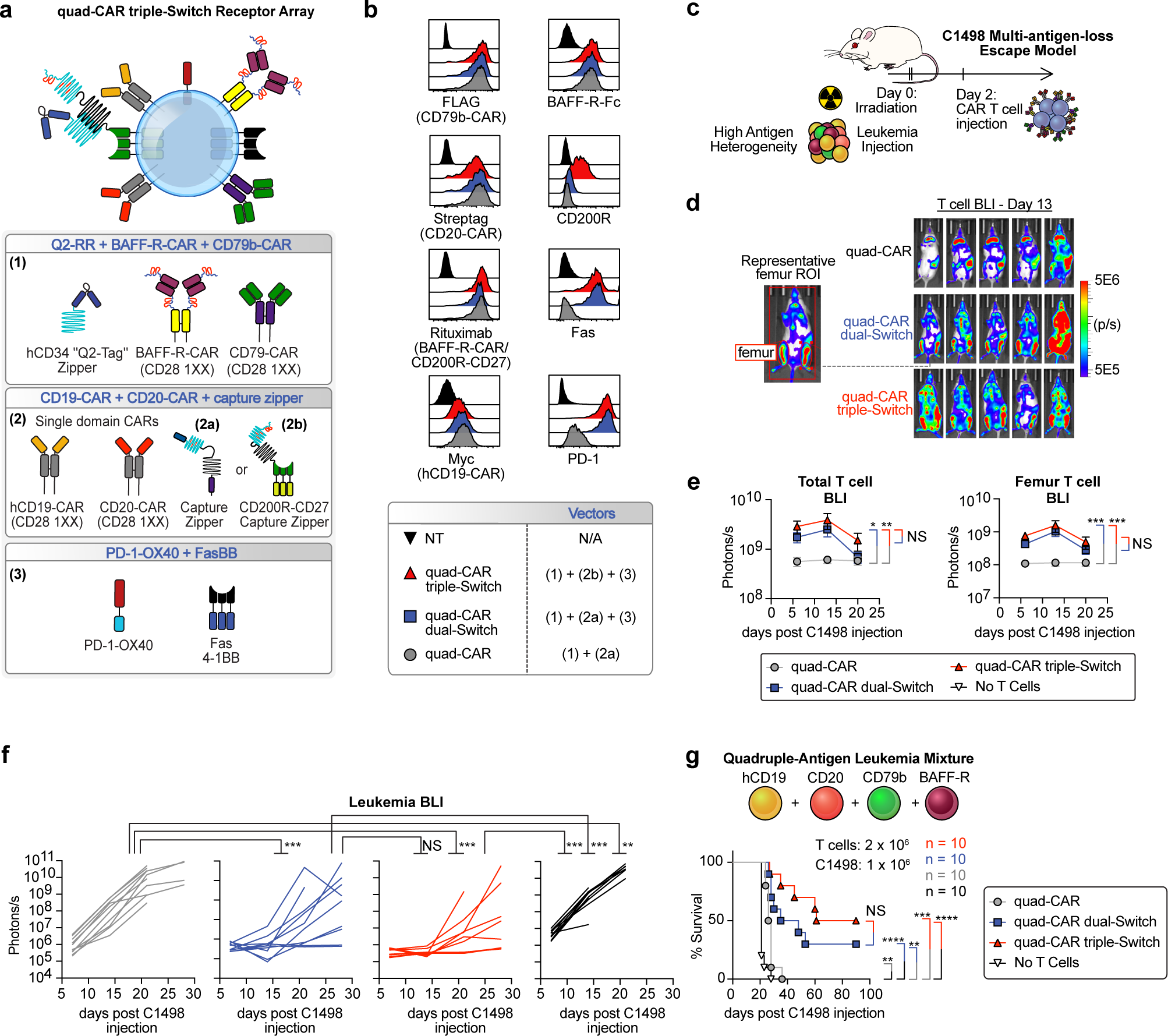
Multi-Switch receptor arrays enhance the activity of quad-CAR T cells. **a**, Diagram depicting quad-CAR dual-Switch and quad-CAR triple-Switch receptor arrays (see Extended Data Fig. 9a for vector maps). **b**, Flow cytometry analysis of CAR and switch-receptor expression in Zip-sorted albino B6 T cells. **c-g**. Sublethally irradiated albino B6 mice were injected with 1:1:1:1 ratio of C1498 singly expressing (hCD19, CD20, CD79b, BAFF-R) and treated with dasatinib-cultured quad-CAR albino B6 T cells co-transduced with Gaussia luciferase vector gLuc-PD-1H-CD24-GPI-P2A-EGFP and expressing switch receptors as depicted. **c**, Experimental setup diagram. **d**, Day 13 T cell BLI and representative example of femur BLI region of interest. **e**, Total T cell BLI kinetics (left) and femur BLI kinetics (right) from two combined experiments. **f**, C1498 BLI from two combined experiments. **g**, Survival. Log-transformed BLI AUC values were compared using a Vardi test with FDR correction. Survival differences were compared via pairwise log-rank test, with FDR correction. *p≤0.05, **p≤0.01, ***p≤0.001, ****p≤0.0001, NS p>0.05.

## Discussion

Tumor antigen-loss escape, lack of T cell persistence, and impairment of T cell function are three barriers that limit achievement of durable disease remissions following CAR T cell therapy^3–5^. Impairment of T cell activity can result from intrinsic T cell exhaustion mechanisms and from extrinsic immune suppression imposed by the tumor microenvironment. By co-expressing multiple CARs and switch receptors that reverse suppressive extracellular signals to positively costimulate cells, we generated CAR T cells able to prevent multi-antigen-loss escape. Importantly, this was achieved while maintaining CAR T cell activity in the setting of antigenically diverse leukemia cells expressing multiple inhibitory checkpoint ligands. To facilitate manufacturing of these complex multi-receptor T cell products, we devised a leucine zipper-based sorting system, which demonstrated great versatility, enabling the selective purification of dual-transduced multi-CAR T cells by magnetically sorting on zipper-tagged safety-switch, dominant negative, and switch receptor molecules.

In multi-CAR T cells, where the Zip-sorting system facilitated selection of T cells with high expression of three or four CARs simultaneously, provision of TNFRSF costimulation through multi-Switch receptor arrays enhanced or was necessary for anti-leukemia activity. We designed all CARs in these arrays in a CD28 CD3ζ 1XX configuration to potentially minimize CAR signaling-induced T cell dysfunction^26^ and to provide the kinetically enhanced anti-tumor activity observed with CD28 costimulation^48^. We incorporated multi-Switch receptor arrays to provide the enhanced proliferation, persistence, and metabolic fitness conferred by TNFRSF signaling^23,47–49^. This configuration that combines CD28-costimulated CARs with TNFRSF-based multi-Switch receptor arrays was highly active in our models, and may provide a simplified configuration for developing functional multi-CAR arrays, which have previously required significant optimization of costimulatory domains^9,59^. This multi-CAR multi-Switch configuration may also facilitate design of CAR T cells with activity against antigenically heterogeneous malignancies such as acute myeloid leukemia^8^ and solid tumors^7^, which can also overexpress inhibitory ligands such as FasL, PD-L1, and CD200^7,60,61^.

In conclusion, we have developed a cell sorting methodology that enables production of T cells incorporating multiple transgenes at high purity. Single-step immunomagnetic selection of dual-vector transduced T cells may also facilitate production of recently described synthetic biology approaches including the logic-gated LINK-CAR^62^ and synNotch^63^ technologies and synZiFTR^64^ synthetic gene circuits that use two vectors to encode necessary components. We utilized the Zip-sorting system to optimize multi-CAR T cell configurations with co-expression of multiple switch receptors. Together, these designs enhanced T cell activity against antigenically diverse targets, providing a unified strategy to overcome tumor immune evasion due to target antigen heterogeneity and expression of multiple inhibitory ligands.

## Methods

### Cell Lines and Culture

BM185^39^ was a gift from Donald Kohn (UCLA). C1498 and Phoenix-Eco were obtained from ATCC. Phoenix-Eco was grown in DMEM (Gibco 11-965-118) supplemented with 10% FCS (Sigma, F2442), GlutaMAX 1x (Gibco, 35050061), and Penicillin-Streptomycin 100 IU/mL / 100 μg/mL (Gibco, 15140-163). C1498 and BM185 cell lines and primary mouse T cells were grown in RPMI 1640 (Gibco, 11-875-119) containing 10% FCS (Sigma, F2442), GlutaMAX 1x (Gibco, 35050061), Penicillin-Streptomycin 100 IU/mL / 100 μg/mL (Gibco, 15140-163), MEM Non-Essential Amino Acids Solution 1x (Gibco, 11140050), Sodium Pyruvate 1 mM (Gibco, 11360070), and 2-mercaptoethanol 55 μM (Gibco, 21985023) (T cell media). T cells were cultured in 50 IU/mL recombinant human IL-2 (rhIL-2) (Proleukin) and in some experiments 1 μM dasatinib (Cayman Chemical, 11498) or 10 mM NAC (Sigma, A9165-25G).

### Vector and DNA Construct Design

Constructs in this study were expressed from retroviruses or transposons. See Supplementary Table 2 for construct sequences and expression vectors used for each construct and see Supplementary Table 3 for amino acid and DNA sequences of specific construct elements present in constructs described in Supplementary Table 2. Vector construct maps are presented in supplemental figures and complete amino acid sequences for all constructs are provided in Supplementary Table 2. CAR and other receptor constructs used murine protein sequences except for Fas 4-1BB, which utilized the human 4-1BB costimulatory domain given greater activity in mice^65^. DNA constructs were generated via standard molecular biology techniques including overlap extension PCR and restriction site-based cloning and ligated directly into pENTR1a no ccdB (Addgene 17398) or the MMLV-based retroviral vector LZRS-Rfa (Addgene 31601). SnapGene 6.2 (SnapGene) was used to design vectors. DNA templates were obtained as gBlocks or gene syntheses from Integrated DNA Technologies (IDT), or purchased from Origene, Addgene, and Sino Biological (Supplementary Table 3). pENTR1a-based constructs were transferred via Gateway cloning (Gateway LR Clonase II Enzyme mix, Invitrogen, 11791020) to LZRS-Rfa, piggybac transposons, or piggybac transposon ITR-flanked retroviral vectors containing attR recombination sites, designed in this study. These vectors were designed for stable high-level expression of retroviral vectors integrated into packaging lines using piggybac transposition. The vectors, named PB-MMLV-puro, PB-MPSV-puro, and PB-SIN-puro include the backbone and ITR and insulator domains of the piggybac transposon vector PB-EF1α-MCS-IRES-GFP (System Biosciences PB530A-2) and contain a 5’ compound SV40 enhancer with hybrid RSV-MMSV LTR (based on SERS11 design^66^), leader sequence containing a modified MESV packaging signal (from MP71^67^), attR sites flanking ccdB and chloramphenicol resistance genes, woodchuck hepatitis virus post-transcriptional regulatory element (WPRE), and followed by 3’ LTR regions from LZRS (MMLV), MP71 (MPSV), or inactivated MMLV LTR from pSIN^68^ (SIN). These vectors also contain an hPGK-promoter driving a Thy1.2-T2A-puroR selection cassette. PB-SFG5.3-BlastR contains the full 5’ LTR and packaging signal from SFG^69^, attR sites flanking ccdB and chloramphenicol resistance genes, WPRE, 3’ SFG LTR, and hPGK-promoter driving a Thy1.2-T2A-BlastR selection cassette. The transposon PB-EF-1α-intron was generated by replacing the EF-1α core promoter in PB-EF1α-MCS-IRES-GFP with the full EF-1α intron-containing promoter. PB-EF-1α-intron-WPRE contains an added 3’ WPRE. gLuc-PD-1H-CD24-GPI-P2A-EGFP encodes a surface-expressed Gaussia luciferase with PD-1 hinge and membrane anchor based on mouse CD24 GPI transfer signal to localize BLI signal to gLuc-expressing cells and also encodes an EGFP reporter. CBR-P2A-hCD8-T2A-puroR is a construct containing click beetle red luciferase (CBR) along with a human CD8 surface reporter and encoding puromycin resistance. PB-SIN-puro BFP-SV40-NFAT-dEGFP, PB-SIN-puro BFP-SV40-AP-1-dEGFP, and PB-SIN-puro BFP-SV40-NFkB-dEGFP are transcription factor reporter vectors with construct elements reported in Supplementary Table 3. PB-EF-1α-intron hCD8-T2A-integrin-alpha-V and PB-EF-1α-intron mCD4-P2A-integrin-beta-3 encode components of human vitronectin receptor and hCD8 and mCD4 reporters, respectively.

### Retrovirus Production

Retroviruses were produced in Phoenix-Eco or Phoenix-Eco alpha-V beta-3, designed to enhance adhesion of Phoenix-Eco to culture flasks. Phoenix-Eco or Phoenix-Eco alpha-V beta-3 were transfected with LZRS-vector constructs using Effectene transfection reagent (Qiagen, 301425) as follows: complexes were generated using 2 μg vector DNA in 16 μL of enhancer, 20 μL of Effectene and used to transfect Phoenix cells plated on a 10 cm dish at 2.5×10^6^ cells one day prior to transfection. On the day following transfection, cells were removed from dishes and assessed by flow cytometry for construct expression and selected in puromycin 2 μg/mL (Santa Cruz Biotech, sc-108071B). Phoenix-Eco or Phoenix-Eco alpha-V beta-3 were transfected with PB-vector constructs as above, with addition of 1 μg of the hyperactive piggyBac transposase vector pCMV-hyPBase^70^ (Sanger Institute, UK) for stable integration. Transfected cells were selected one day after transfection with puromycin 2 μg/mL (Santa Cruz Biotech, sc-108071B) or blasticidin 10 μg/mL (Santa Cruz Biotechnology, sc-204655). Retroviral supernatant was collected from fully selected stable packaging lines grown from T175 flasks. Supernatants were filtered through 50 mL syringes fitted with Millex-HV Syringe Filters (Millipore, 0.45 µm, PVDF, SLHVR33RS) and polyethylene glycol solution concentrate was added (5X concentrate: PEG 8000 MW Promega, V3011, 40% weight/volume containing 2.4% NaCl weight/volume), and virus was precipitated over 1-2 days at 4C. Precipitated virus was centrifuged at 3000xg for 15 minutes at 4C and pellets were resuspended in 500 µL of T cell media and frozen at -80C or used directly.

### Retroviral Transduction

Tumor cell lines were transduced with retroviral supernatant containing polybrene 8 μg/mL (Sigma, H9268-10G) in 24-well plates by spinfection at 1500 RPM at 32C for 1 hour. Cells were transferred to T25 flasks on the following day for expansion. For primary mouse T cell transduction, splenic T cells were enriched by negative selection of splenocytes with anti-CD19 microbeads to deplete B cells (Miltenyi, 130-121-301) and were stimulated on plate-bound anti-CD3/CD28 for 1 day (2 μg/mL each, clones 145-2C11 and 37.51, respectively, BioXCell). Activated T cells were transduced on day 1 after stimulation using combinations of PEG-precipitated retroviral concentrates encoding different constructs adsorbed onto non-tissue culture treated 6-well plates coated with anti-CD3/CD28 2 μg/mL each and retronectin 20 μg/mL (Takara, T100B) at 1-2×10^6^ cells/well. On day 2, T cells were either re-transduced, or transferred to 6-well plates coated with anti-CD3/CD28. Cells were removed on day 3, Zip-sorted, and expanded in 50 IU/mL rhIL-2 (Proleukin).

### Target Cell Line Construction

BM185-CD19 was constructed by transducing BM185 with LZRS ffluc-Thy1.1-Neo and CD38 was deleted for use in concurrent studies using Cas9-NLS (UC Berkeley MacroLab), CD38 gRNA UAAAUUCAUAGUUAGCCAUU (Synthego), and Lonza SF buffer kit (Lonza, V4XC-2032) with a 4D Nucleofector using code DN100. Subclones were identified that were CD19^+^ CD38^-^. BM185-CD20 was similarly generated as a clone uniformly expressing the LZRS CD20 and LZRS ffluc-Thy1.1-Neo transgenes. CD19 was deleted with CD19 gRNA UGAUUCAAACUGCUCCCCCG and CD38 deleted with the CD38 gRNA, after which cells were negatively selected for CD19 (CD19 microbeads, Miltenyi, 130-121-301) and CD38 (anti-CD38 PE, anti-PE microbeads, Miltenyi, 130-048-801). BM185-CD19-CD20 expresses LZRS ffluc-Thy1.1-Neo and was previously described^38^. BM185-CD19-FasL was generated from a BM185-CD19 ffluc-Thy1.1-Neo clone generated for concurrent studies with deletion of CD38 and CD22 (CD22 gRNA UGUCAUUGGCACGUAUCGGG, Synthego) and transduced with LZRS FasL and subcloned. BM185-CD20-PD-L1 and BM185-CD20-CD200 were similarly generated from CD38, CD22-deleted clones modified to express PD-L1 (LZRS PD-L1) or CD200 (PB-EF-1α-intron CD200, pCMV-hyPBase, SF buffer kit V4XC-2032, Lonza 4D Nucleofector, code DN100). Transposition reactions used 1-2 μg of vector DNA and 0.5-1.0 μg of pCMV-hyPBase transposase DNA. These cell lines were then subcloned. C1498-CD19 and C1498-CD20 versions were generated by transduction with SFG CD19 and LZRS CD20 and LZRS ffluc-Thy1.1-Neo and immunomagnetically sorted for high transgene expression. However, long-term retroviral expression in C1498 was unstable, so we generated a stable C1498 CBR-hCD8-puro clone using PB-EF-1α-intron transposon and generated subcloned versions individually transposed with the PB-EF-1α-intron transposon vectors encoding CD19, hCD19, CD20, CD79b1, or BAFF-R (transposon, pCMV-hyPBase, SF buffer kit V4XC-2032, Lonza 4D Nucleofector, code DA100). CD79bΔ refers to a chimeric protein comprising mouse CD79b extracellular domain with a CD28TM and truncated CD3ζΔ to promote surface expression in the absence of CD79a. For Incucyte live-cell microscopy analysis, target cells were transduced with PB-MSPV-puro iRFP713-P2A-hygro-E2A-TAA-WPRE and selected in hygromycin B (Santa Cruz Biotechnology, sc-29067) at 0.8 mg/mL to enable near-infrared imaging.

### Zip-Sorting

Transduced C1498 or primary T cells were incubated with anti-FLAG (Miltenyi, 130-101-591) or anti-CD34 (Miltenyi, 130-046-702) beads at 30 μL of beads per 10^7^ cells for 30 minutes at 4C in PBS 2 mM EDTA + 0.5% BSA, washed in PBS 2 mM EDTA + 0.5% BSA, centrifuged 1200 RPM x 5 minutes and resuspended in 500 μL PBS 2 mM EDTA + 0.5% BSA. Cells were sorted on LS columns (Miltenyi, 130-042-401) by washing 3x (1 mL, 2 mL, 3 mL) with PBS 2 mM EDTA + 0.5% BSA and eluting with 5 mL T cell media. Zip-sorted T cells were used for in vitro experiments on days 4-6 post-stimulation and injected into mice for in vivo experiments on day 5 post-stimulation. Sort yield is defined as 100*(dual-transduced cells recovered/dual-transduced cells present pre-sort).

### Continuous Live-Cell Microscopy

T cells were transduced with either PB-SFG5.3-BlastR EGFP (constitutive EGFP) or NFκB reporter vectors (inducible, destabilized EGFP reporter), added at 2 x10^4^ cells/well to 96-well flat-bottom plates, and co-cultured with iRFP713-expressing target lines at varying E:T ratios in T cell media without rhIL-2. T cell and tumor cell line fluorescence was imaged simultaneously every 3 hours with 10x objective, 4 images per well, in an Incucyte SX5 (Sartorius). For stress-test experiments, targets were added back to the plate at 1:1 initial E:T ratio at 24 and 48 hours after initial assay setup. For T cell cluster analysis, green fluorescence object threshold was set to minimum of 5000 μm^2^, edge split off, eccentricity maximum 1.0, hole fill 0 μm^2^. Data were analyzed using Incucyte Software v2021A.

### Bioluminescence-based Target Lysis Assay

T cells were incubated in triplicate in 96-well U bottom plates for 24 hours at varying E:T ratios with 0.5-1×10^4^ luciferase-expressing targets in T cell media lacking rhIL-2. A no-T cell row was added to obtain relative maximum luciferase activity. For experimental readout, luciferin (Gold Bio LUCK-2G) was added to wells to achieve final 200 μg/mL concentration and luciferase activity was analyzed on a Tecan SPARK luminometer/fluorimeter (Tecan). Target relative percent viable values were calculated as 100*(experimental well activity units / maximum activity units).

### Flow Cytometry

See Supplementary Table 4 for antibodies used. Flow cytometry analysis acquisition was performed on an LSR-II or FACSymphony X50 using FACSDiva software (BD Biosciences). Analysis was performed using FlowJo software (BD Biosciences, version 10.8.1). Cell viability was assessed with DAPI (Calbiochem, 5087410001). MFI refers to geometric mean fluorescence intensity. Intracellular flow cytometry analysis was performed by first antibody staining cells for surface markers and live/dead status using LIVE/DEAD Fixable Blue Dead Cell Stain Kit (Invitrogen, L23105), followed by permeabilization using the Foxp3 / Transcription Factor Staining Buffer Set (Invitrogen, 00-5523-00) and antibody stained for intracellular contents. CAR expression was detected by flow cytometry with antibodies against affinity tags including Myc, Streptag, FLAG, hCD20 mimotope (Rituximab-APC), and the hCD34 tandem epitope^35^ with their respective antibodies. Streptag was also detected using Streptactin-PE for some experiments (Iba Biosciences, 6-5000-001). Rituximab was obtained from the MSKCC pharmacy and APC conjugated (APC Conjugation Kit - Lightning-Link, Abcam, ab201807). Anti-streptag purified antibody (Genscript) was also APC conjugated. The BAFF-R CAR was stained with 1 μg of hBAFF-R hFc (Sino Biological, 16079-H02H), followed by anti-hFc antibody. Similarly, PD-1-DNR interaction with PD-L1 was assessed by staining with 1 μg Recombinant Mouse PD-L1/B7-H1 Fc Chimera Protein (R&D Systems, 1019-B7-100), followed by anti-hFc antibody.

### Complement Lysis Assay

T cells were incubated at 5×10^4^ cells/well in 96-well U bottom plates with rabbit complement (final concentration 10%, Cedarlane, CL3051) ± 100 μg/mL anti-Thy1.1 (clone 9E12, BioXCell) for 30 minutes at 37C. Subsequently, viable cells were enumerated by flow cytometry, measuring DAPI-negative viable cells using CountBright Beads (Invitrogen, C36950). Relative T cell survival was calculated as 100*(viable cells: antibody + complement / viable cells: complement only).

### iCaspase9 Activation Assay

For dimerizer titration, T cells were incubated at 3×10^4^ cells/well in 96-well U bottom plates for 24 hours in T cell media containing 50 IU/mL rhIL-2 and varying concentrations of AP20187 (B/B homodimerizer, Takara, 635058). After 24 hours, cells were analyzed for DAPI-negative viable cells using CountBright Beads (Invitrogen, C36950). Relative T cell survival was calculated as 100*(viable cells: dimerizer / viable cells: DMSO).

### Reactive Oxygen Species Assessment

Unstimulated day 5 T cells were plated at 5×10^4^ cells/well in 96-well U bottom plates, washed once with 200 μL of PBS, resuspended in 100 μL of PBS containing 1 μM CM-H_2_-DCFDA (Invitrogen, C6827) or 5 μM Mitosox Red (Invitrogen, M36008) for 30 minutes at 37C. Cells were washed twice in 200 μL of PBS + 0.5% BSA, stained with anti-CD4/CD8, and analyzed by flow cytometry for DAPI-negative viable cells.

### Transcription Factor Reporter Assay

T cells were transduced with transcription factor reporter vectors on day 1 post-stimulation (to ensure equivalent reporter transduction among T cells subsequently transduced with different constructs) and on day 2 the cells were transduced with different CAR construct vectors. T cells were subsequently Zip-sorted or left unsorted for further analysis. T cells were assessed for EGFP induction by flow cytometry following culture with or without target cells in BFP-transduction reporter-positive cells.

### In vivo experiments

Female BALB/cJ (Jackson Laboratory, 00651) and B6(Cg)-*Tyr^c-2J^*/J albino B6 (Jackson Laboratory, 000058) mice were purchased at 6-8 weeks of age and used in experiments at 7-12 weeks of age. Albino B6 were utilized to enhance BLI sensitivity given absence of fur pigmentation. CD45.1 congenic BALB/c mice (Jackson Laboratory, 006584) were bred at MSKCC. Animal studies were conducted in the MSKCC vivarium under a protocol approved by the MSKCC Institutional Animal Care and Use Committee. Mice were evaluated at least twice daily and euthanized when reaching any of the following humane endpoints: weight loss > 25%, labored breathing, moribund status, hind-limb paralysis, development of ascites, tumor > 2 cm or interfering with bodily functions.

### BM185 pre-B Acute Lymphoblastic Leukemia Mouse Model

BALBc/J mice were sublethally irradiated with 450 cGy of gamma radiation (Gammacell, cesium source), rested for 4 hours, and then injected with varying doses of BM185 cell lines via tail vein in 200 μL of DMEM without additives (day 0). Mice were randomized into groups following leukemia injection. On day 2, Zip-sorted T cells were injected into the retroorbital plexus in 150 μL of DMEM. T cell and BM185 cell doses are depicted in figures above survival or BLI plots. Mice were evaluated daily for evidence of reaching humane endpoints described in the *In vivo* experiments section. Mice were serially assessed for leukemia progression via firefly luciferase (ffluc)-based BLI. In some experiments, spleens or bone marrow (BM) were obtained at the time of euthanasia for further analysis. Spleens and BM were dissociated through 40-micron filters, red blood cell lysed (Hybri-Max, Sigma, R7757), and stained with anti-CD3ε, Thy1.1, CD19, and CD20. Spleens or BM with < 0.1% Thy1.1^+^ CD3^-^ BM185 cells or < 10 events were excluded from analysis of BM185 surface phenotype. In some experiments, mice were bled via retroorbital plexus, blood was red blood cell-lysed, and stained for flow cytometry analysis.

### C1498 Acute Myeloid Leukemia Mouse Model

Albino B6 mice were sublethally irradiated with 550 cGy of gamma radiation (Gammacell, cesium source), rested for 4 hours, and then injected with varying doses of C1498 cell lines via tail vein in 200 μL of DMEM without additives (day 0). Mice were randomized into groups following leukemia injection. On day 2, Zip-sorted T cells were injected into the retroorbital plexus in 150 μL of DMEM. T cell and C1498 cell doses are depicted in figures above survival or BLI plots. Mice were evaluated daily for evidence of reaching humane endpoints described in the *In vivo* experiments section. Mice were serially assessed for leukemia progression via CBR luciferase-based BLI. In some experiments, BM was obtained at the time of euthanasia for further analysis. BM was dissociated through 40-micron filters, red blood cell lysed (Hybri-Max, Sigma, R7757), and stained for hCD8 and tumor target antigens. BM with < 0.1% hCD8^+^ C1498 cells or < 10 events was excluded from C1498 surface phenotype analysis. In BM of some CAR T cell treated mice, we observed an amorphous debris that simultaneously stained positively for all flow markers: hCD8, CD19, CD20, CD79b, and BAFF-R. This population was gated out of analyses. The C1498 model was less predictable than BM185, with mice sometimes dying overnight despite looking otherwise healthy on the prior night. Additionally, a subset of CAR T cell treated mice apparently cleared leukemia from the BM with extramedullary progression. Therefore, the number of available BM samples with C1498 to assess for antigen-loss escape was diminished compared with the BM185 model.

### Bioluminescence Imaging (BLI)

For serial quantitative assessment of leukemia, mice were injected with D-luciferin (Gold Bio, LUCK-2G) at 150 mg/kg dose intraperitoneally. Ten minutes after injection, isoflurane-anesthetized mice were imaged using an IVIS Spectrum CT imaging system (PerkinElmer). For serial quantitative assessment of T cells, mice were injected with 100 μg of water-soluble Coelenterazine (NanoLight Technology, 3031) into the retroorbital plexus and imaged immediately. Mice were imaged individually following injection.

### Statistical Analysis

For BLI curves, groups were compared using area under the curve (AUC) analysis performed on log-transformed BLI values using a Vardi test^71^ with the function aucVardiTest in the R package clinfun^72^. False discovery rate (FDR) correction was applied to account for multiple comparisons. Pairwise log-rank tests were performed by the function *pairwise_survdiff* in the R package *survminer*^73^, followed by FDR correction for multiple tests. All other statistical tests were performed using GraphPad Prism, with test type described in figure legends.

## Supporting information

Table S2 - construct sequences

Table S3 - element sequences

Table S4 - antibodies

## Data Availability

The data generated in this study are available upon request from the corresponding author.

## Acknowledgements

This research was supported by National Cancer Institute award numbers R01-CA228358, R01-CA228308, P30 CA008748 MSK Cancer Center Support Grant/Core Grant, and P01-CA023766; National Heart, Lung, and Blood Institute (NHLBI) award number R01-HL123340 and R01-HL147584; National Institute on Aging award number P01-AG052359, and Tri-Institutional Stem Cell Initiative. Additional funding was received from The Lymphoma Foundation, The Susan and Peter Solomon Family Fund, The Solomon Microbiome Nutrition and Cancer Program, Cycle for Survival, Comedy vs. Cancer, Parker Institute for Cancer Immunotherapy, Paula and Rodger Riney Multiple Myeloma Research Initiative, and the Starr Cancer Consortium. S.E. James is supported by a K08 career development award NCI K08-CA252157. S.E. James was also supported by a Young Investigator award from the American Society for Clinical Oncology, an Amy Program Award from the Be the Match Foundation, and a Bridge Scholar award from the Parker Institute for Cancer Immunotherapy. S. Chen was supported by a Research Fellowship from the German Research Foundation (Deutsche Forschungsgemeinschaft, DFG). S. DeWolf receives research support from the MSK Leukemia SPORE Career Enhancement Program and the Gerstner Physician Scholar program. J. U. Peled is supported by an NHLBI NIH Award K08-HL143189 and the MSKCC Cancer Center Core Grant NCI P30 CA008748. S.A. Vardhana is supported by a K08 career development award NCI K08-CA237731 and the Parker Institute for Cancer Immunotherapy. C.A. Klebanoff is supported in part by NIH/NCI R37 CA259177, R01 CA269733, P30 CA008748 and the Parker Institute for Cancer Immunotherapy.

## Authors Contributions

**Conceptualization**: S.E. James, L. Jahn, and M.R.M. van den Brink

**Development of Methodology**: S.E. James, L. Jahn, S.A. Vardhana, C.A. Klebanoff, and M.R.M. van den Brink

**Funding acquisition:** S.E. James and M.R.M. van den Brink

**Investigation:** S.E. James, S. Chen, B.D. Ng, J.S. Fischman, L. Jahn, A.P. Boardman, A. Rajagopalan, H.K. Elias, A. Massa, D. Manuele, K.B. Nichols, A. Lazrak, and N. Lee

**Formal Analysis:** S.E. James, S. Chen, T. Fei, S. DeWolf, and J. U. Peled

**Writing – original draft:** S.E. James

**Writing – review & editing:** S.E. James, S. Chen, B.D. Ng, J.S. Fischman, L. Jahn, A.P. Boardman, A. Rajagopalan, H.K. Elias, S. DeWolf, J. U. Peled, S.A. Vardhana, C.A. Klebanoff, and M.R.M. van den Brink

**Supervision:** M.R.M. van den Brink

---

**Supplementary Table 1:**
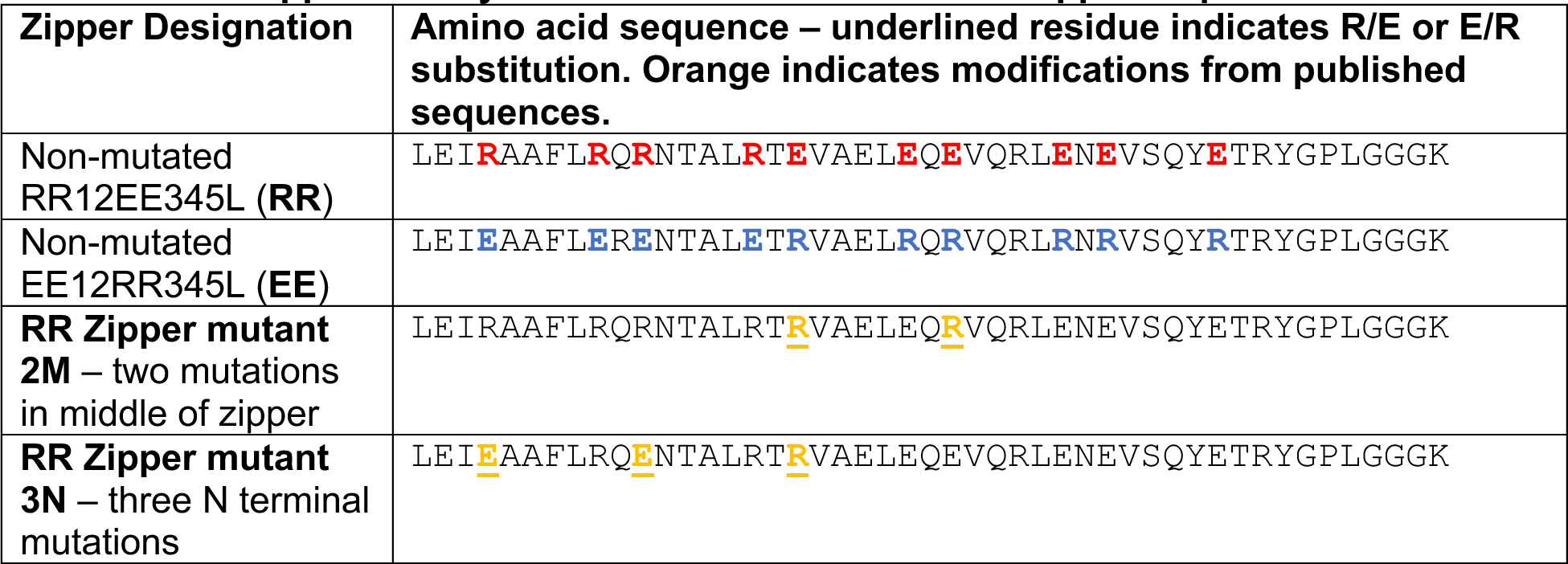
Mutant RR12EE345L zipper sequences.

**Extended Data Fig. 1:**
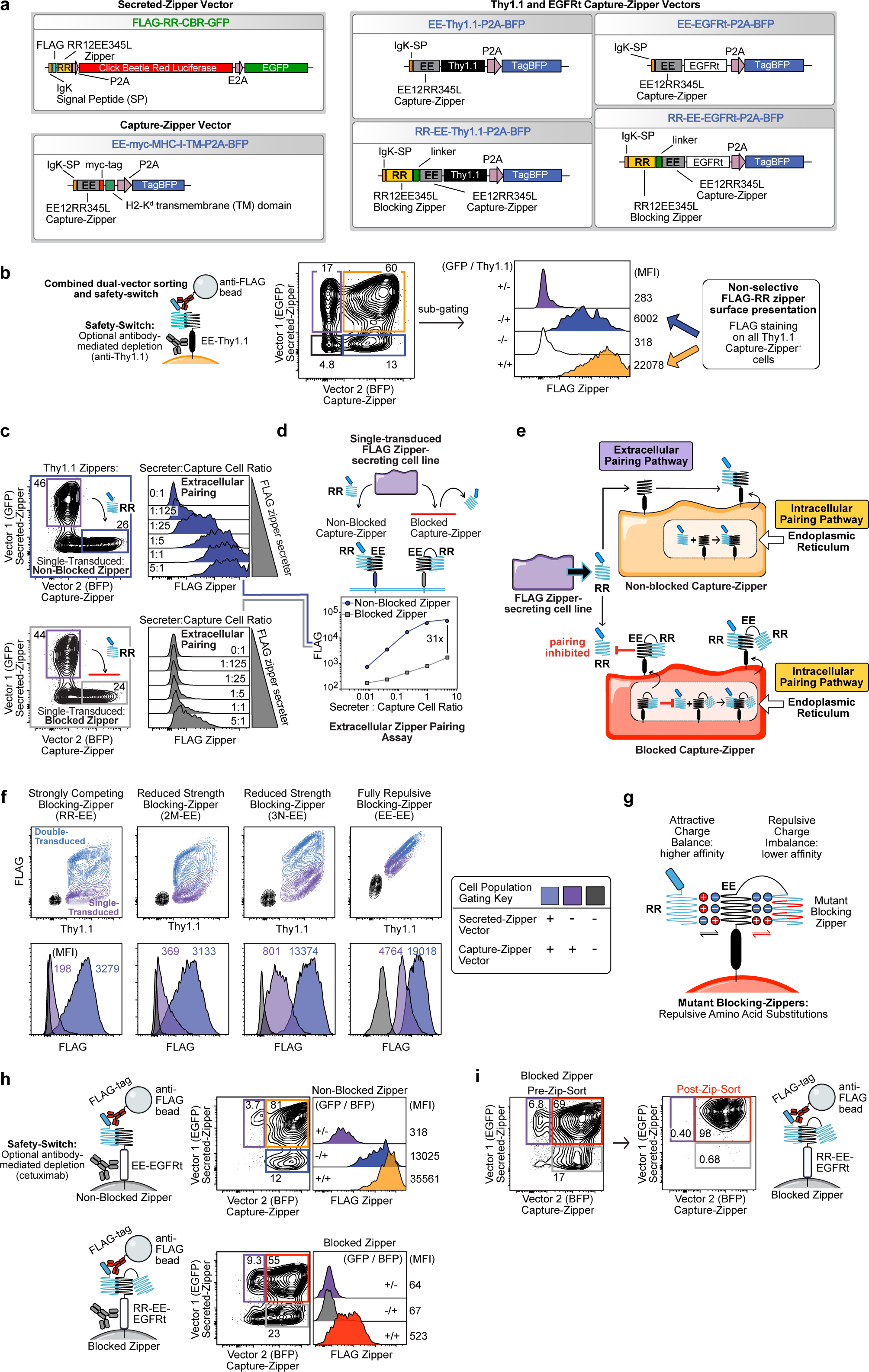
Covalently linked blocking-zipper inhibits extracellular leucine zipper pairing. **a**, Zip-sorting system vector maps. **b**, Flow cytometry analysis of C1498 cells co-transduced with FLAG-RR-CBR-GFP and EE-Thy1.1-P2A-BFP vectors. **c**, Flow cytometry analysis of co-culture of single-transduced FLAG-zipper-secreting FLAG-RR-CBR-GFP C1498 with C1498 cells single-transduced with either (top) EE-Thy1.1-P2A-BFP unblocked or (bottom) RR-EE-Thy1.1-P2A-BFP blocked capture-zipper vectors at depicted cell ratios for 48h. FLAG staining depicts extracellular FLAG-RR zipper pairing on single-transduced capture-zipper^+^ C1498 cells secreted into the media by single-transduced FLAG-RR-secreting C1498 cells. **d**, FLAG-RR zipper surface expression on C1498 cells expressing blocked or non-blocked capture-zippers depicted in panel c. n=1 transduction for each cell line and n=3 co-culture experiments. Data are ±SEM of triplicate samples. Error bars were too small to depict. **e**, Diagram depicting proposed intracellular and extracellular zipper pairing pathways. Non-blocked zippers can obtain secreted FLAG-RR zipper extracellularly from any cell able to secrete it. The blocking-zipper inhibits pairing with secreted FLAG-RR. FLAG-RR pairs with the capture-zipper more strongly intracellularly vs. extracellularly, resulting in greater surface FLAG-RR expression in dual-transduced vs. single-transduced cells. **f,** Flow cytometry contour plots and histograms of mixed populations of C1498 cells single and dual-transduced with FLAG-RR-CBR-GFP and RR-EE-Thy1.1-P2A-BFP vectors with different repulsive mutations in blocking-zippers (See Supplementary Table 1). “EE” blocking-zipper is engineered to be fully repulsive against EE capture-zipper and maximally attractive towards the FLAG-RR zipper. Remaining mutants contain varying numbers of repulsive mutations “2 or 3” in the N-terminal “N” or middle “M” regions of the blocking-zipper. Representative of n=3 separate transductions. **g**, Diagram depicting predicted effect of repulsive amino acid substitution on zipper binding affinity. **h**, Flow cytometry analysis of C1498 cells dual-transduced with FLAG-RR-CBR-GFP and either (top) non-blocked EE-EGFRt-P2A-BFP or (bottom) blocked zipper RR-EE-EGFRt-BFP. **i**, Zip-sort of FLAG-RR-CBR-GFP/RR-EE-EGFRt-P2A-BFP C1498 cells. Representative of n=2 transductions and Zip-sorts.

**Extended Data Fig. 2:**
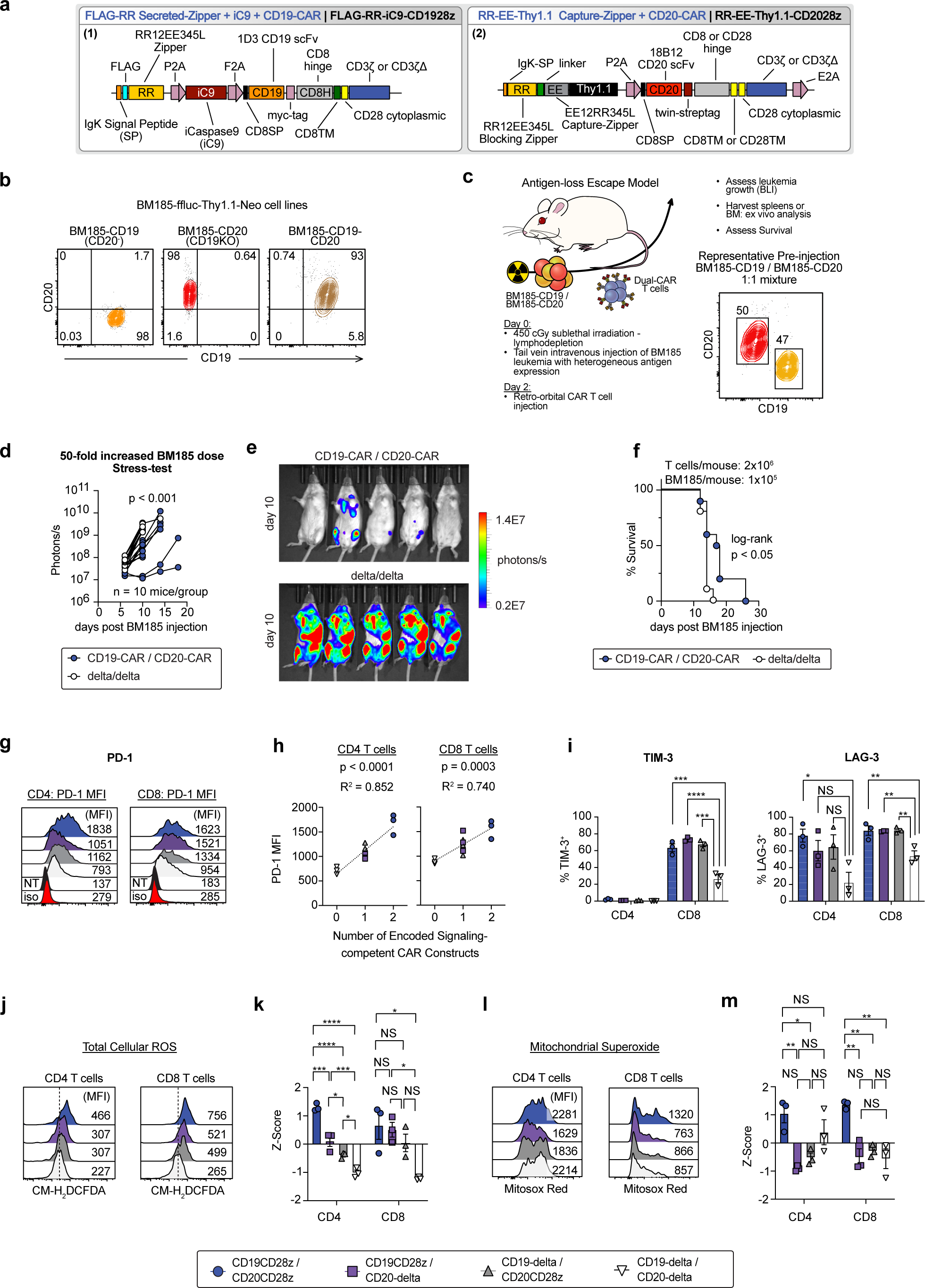
Zip-sorted dual-CAR T cells demonstrate CAR-dose-dependent upregulation of ROS and inhibitory receptor expression. **a**, Maps of vectors for constructs depicted in Fig. 2. **b**, Flow cytometry analysis of BM185-ffluc-Thy1.1-Neo cell lines expressing combinations of CD19 and CD20. **c**, Schematic depicting BM185 syngeneic mouse model of antigen-loss escape in CAR T cell immunotherapy for acute lymphoblastic leukemia. **d-f**. Sublethally irradiated BALB/c mice were injected with 1:1 mixture of BM185-CD19/BM185-CD20 at 1×10^5^ BM185/mouse (50x increased dose vs. Fig. 2g) and treated with Zip-sorted BALB/c CAR T cells on day 2. **d**, Leukemia BLI (ffluc) from two combined experiments. Log-transformed BLI values were compared using a Vardi test to compare AUC values with FDR correction for multiple comparisons. **e**, Day 10 BLI images from representative experiment. **f**, Survival, compared via log-rank test. **g**, PD-1 expression of resting Zip-sorted BALB/c CAR T cells. **h**, Linear regression analysis of unstimulated T cell PD-1 expression vs. number of signaling-competent CARs expressed, n=3 donors. **i**, Immunophenotypic analysis of unstimulated Zip-sorted BALB/c CAR T cells; mean ± SEM from n=3 donors. **j**, Total cellular ROS (CM-H_2_DCFDA) analysis of resting Zip-sorted CAR T cells. **k**, Z-score normalized mean CM-H_2_DCFDA MFI results of n=3 biological replicates. **l**, Mitochondrial superoxide (Mitosox Red) analysis of Zip-sorted CAR T cells. **m**, Z-score normalized mean Mitosox Red MFI results of n=3 biological replicates. Statistical differences for ROS and mitochondrial superoxide were compared using one-way ANOVA, with Tukey’s test for pairwise comparisons. *p≤0.05, **p≤0.01, ***p≤0.001, ****p≤0.0001, NS p>0.05.

**Extended Data Fig. 3:**
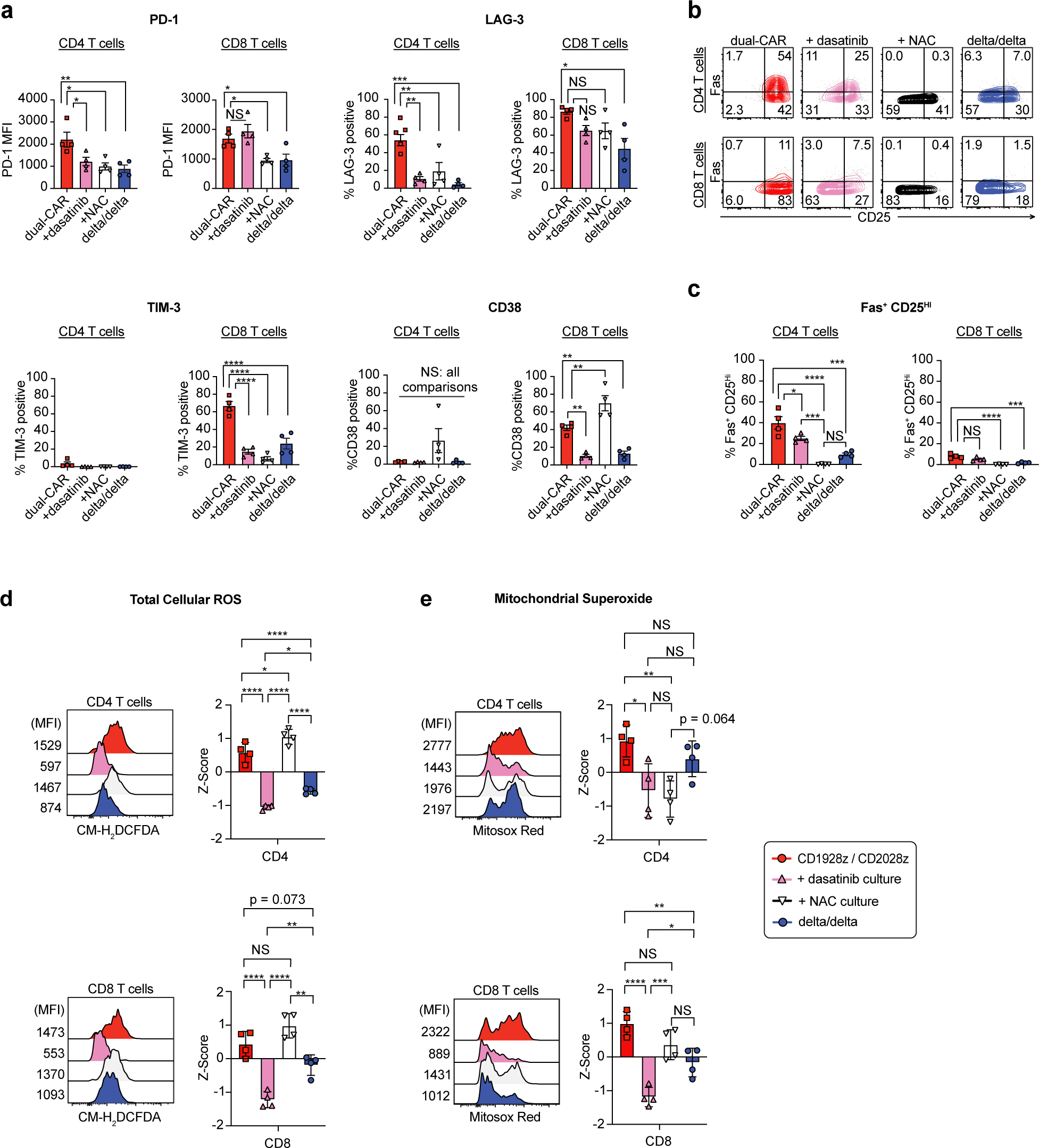
Culture of CAR T cells with NAC and dasatinib inhibits upregulation of inhibitory receptors and ROS. **a-c. a**, Activation/exhaustion marker expression of resting Zip-sorted CD1928z/CD2028z dual-CAR BALB/c T cells cultured with 1 μM dasatinib (2 days), 10 mM NAC (3 days), or DMSO (3 days). **b-c**, Fas, CD25 expression. Data are mean ±SEM of n=4 donors (biological replicates). **d** and **e**, CM-H_2_DCFDA total cellular ROS and Mitosox Red mitochondrial superoxide analysis measurement, respectively, of cells treated as in panel a. (Left) Representative flow cytometry plots. (Right) Z-score normalized MFI means from n=4 replicate experiments with different donors. Statistical differences were compared using one-way ANOVA, with Tukey’s test. *p≤0.05, **p≤0.01, ***p≤0.001, ****p≤0.0001, NS p>0.05.

**Extended Data Fig. 4:**
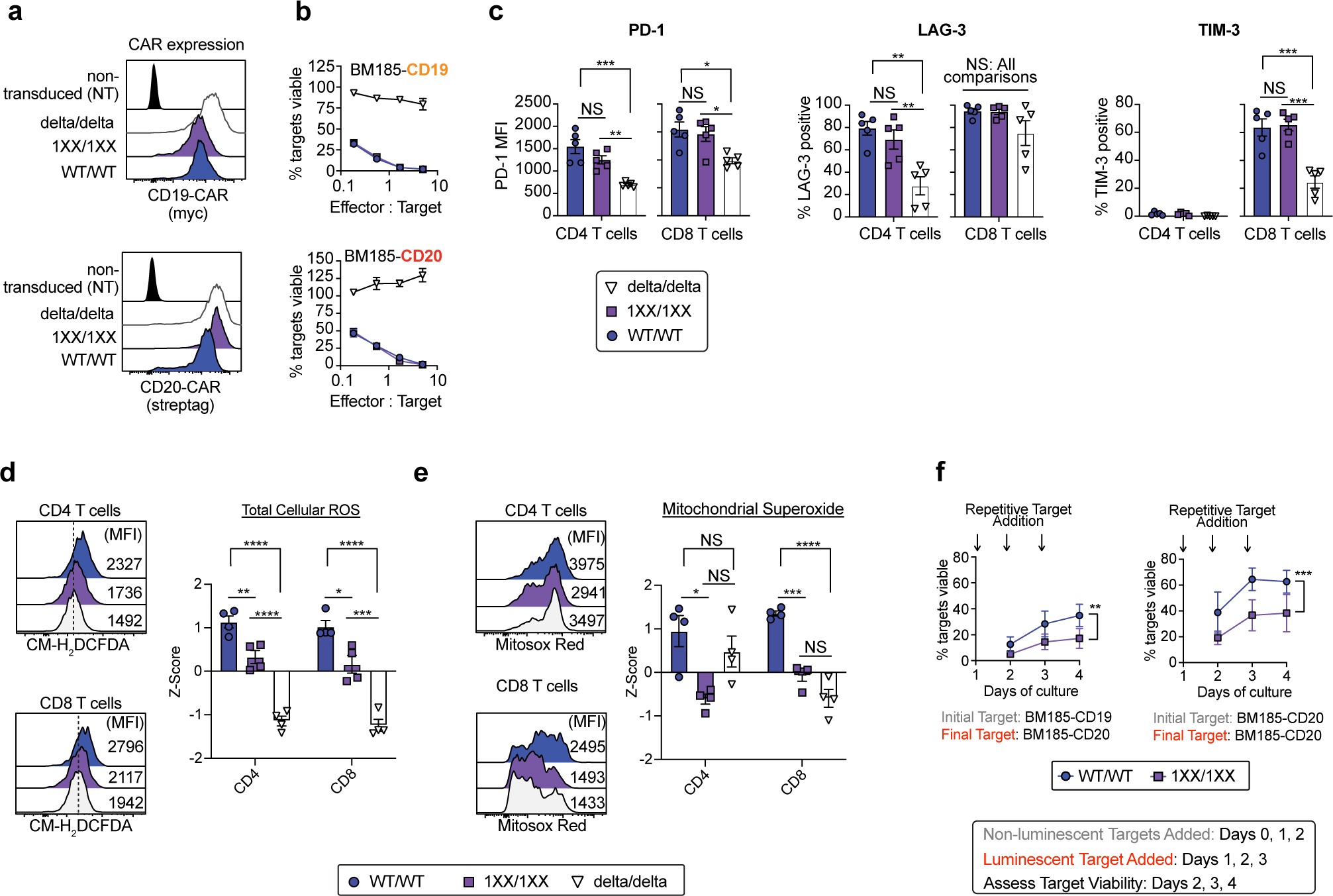
CD3ζ ITAM mutant 1XX dual-CAR T cells demonstrate reduced tonic ROS production and retain greater target lysis capacity following repetitive target encounter. **a**, Flow cytometry analysis of CAR expression in Zip-sorted CD19/CD20 dual-CAR or non-transduced (NT) BALB/c T cells as depicted. **b**, 24h luciferase-based target lysis assay with Zip-sorted CD19/CD20 dual-CAR BALB/c T cells as depicted. Data are mean ±SEM for triplicate wells, from n=2 donor replicate experiments. **c**, Flow cytometry analysis of activation/exhaustion expression on unstimulated cultured Zip-sorted dual-CAR BALB/c T cells. Data are mean ±SEM for n=5 donor T cell transductions. **d-e**, CM-H_2_DCFDA total cellular ROS and Mitosox Red mitochondrial superoxide analysis measurement, respectively, of cells described in panel c. (Left) Representative flow cytometry. (Right) Z-score normalized MFI means from n=4 replicate experiments with different donors. Statistical differences in immunophenotype, ROS, and mitochondrial superoxide were analyzed using one-way ANOVA, with Tukey’s test. **f**, Luciferase-based serial target challenge “stress-test” assay where Zip-sorted CD19/CD20 dual-CAR BALB/c T cells are first challenged with luciferase-negative targets (E:T = 2:1) for varying numbers of daily target additions, followed by a final diagnostic luciferase-positive target addition for 24h for read out of residual target lysis capacity. Data are mean ±SEM from n=3-4 means (biological replicates) of independent experiments. Target viability AUC values were calculated and compared in Prism and compared via a two-tailed t-test. *p≤0.05, **p≤0.01, ***p≤0.001, ****p≤0.0001, NS p>0.05.

**Extended Data Fig. 5:**
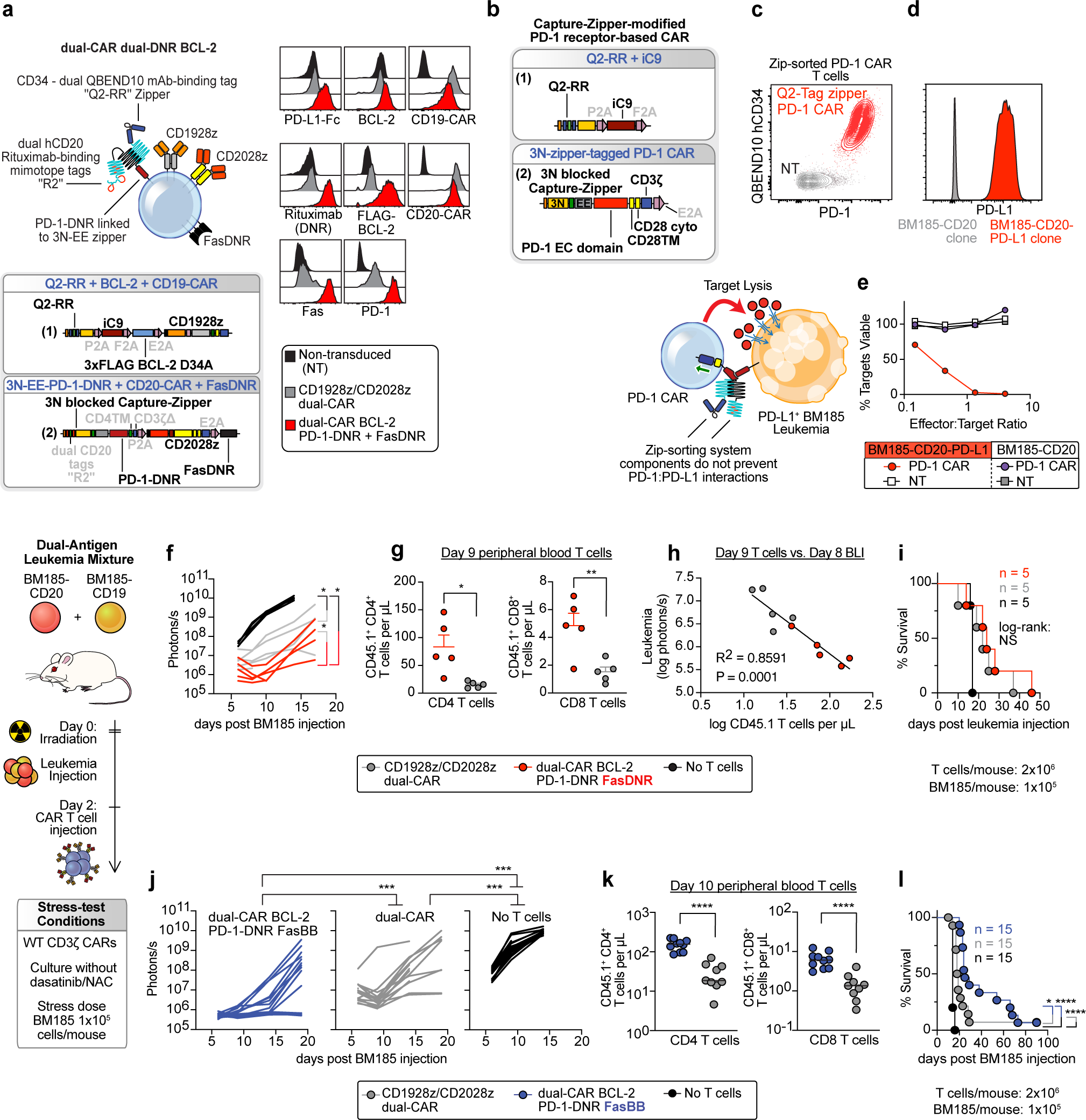
Co-expression of BCL-2 with dominant negative receptors or combination with a single switch receptor transiently enhances anti-leukemia activity of dual-CAR T cells. **a**, (Left) Diagram and maps for vector set encoding Zip-sorting system, dual-CAR, BCL-2 D34A caspase-cleavage resistant mutant, iC9, 3N-mutant blocked zipper-tagged PD-1-DNR and FasDNR (DNR; dominant negative receptor). 3N mutant was used to increase affinity-tag zipper surface expression (See Extended Data Fig. 1f). (Right) Flow cytometry analysis of Zip-sorted BALB/c T cells dual-transduced with this vector set. **b**, Maps for vector set encoding Q2-RR-iC9 and 3N zipper-tagged PD-1-CAR. **c**, Expression of PD-1-CAR and surface bound Q2-RR zipper. **d**, PD-L1 expression on BM185-CD20-PD-L1 clone. **e**, Luciferase-based 24h target lysis assay using Zip-sorted Q2-RR-iC9/3N-EE-PD-1-CAR BALB/c T cells or non-transduced T cells. Data are mean ± SEM of triplicate wells for a representative experiment from n=2 donor replicates. **f-i**. Sublethally irradiated BALB/c mice were injected with a 1:1 mixture of BM185-CD19 and BM185-CD20 and treated with Zip-sorted CD45.1^+^ congenic BALB/c T cells. **f**, Leukemia BLI (ffluc) from single experiment. **g**, Flow cytometry analysis of CAR T cells in peripheral blood on day 9. **h**, Linear regression analysis of leukemia BLI signal vs. blood CD45.1^+^ T cell concentration. **i**, Survival. **j-l**. Experimental setup as in panel f, but mice were treated with dual-CAR T cells incorporating FasBB switch receptor instead of FasDNR. **j**, Leukemia BLI from three combined experiments. **k**, Flow cytometry analysis of CAR T cells in peripheral blood on day 10, combined from two experiments. **l**, Survival from three combined experiments. Log-transformed BLI AUC values were compared using a Vardi test with FDR correction. Blood T cell counts were compared with a two-tailed t-test. Survival was compared with log-rank or pair-wise log-rank comparison with FDR correction. *p≤0.05, **p≤0.01, ***p≤0.001, ****p≤0.0001, NS p>0.05.

**Extended Data Fig. 6:**
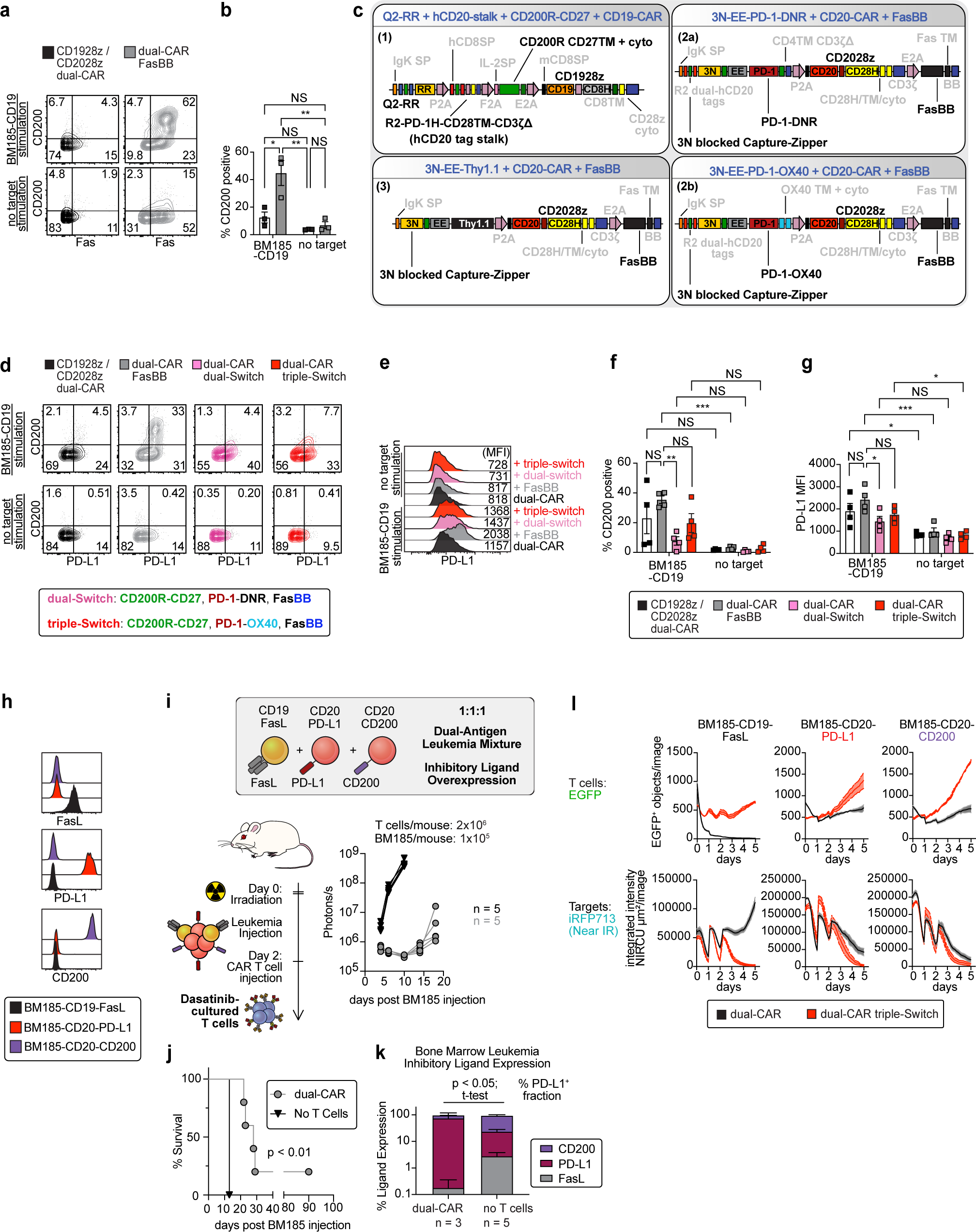
Multi-Switch receptor arrays enhance the proliferation and anti-leukemia activity of dual-CAR T cells. **a-b**, Flow cytometry analysis of Zip-sorted dual-CAR BALB/c T cells ± FasBB switch receptor co-cultured for 24h with BM185-CD19 (E:T = 1:1) or left unstimulated. Data are mean ± SEM (biological replicates) of triplicate wells from n=3 donor experiments. **c**, Vector maps for dual-CAR dual-Switch PD-1-DNR (CD200R-CD27, PD-1-DNR, FasBB), dual-CAR triple Switch (CD200R-CD27, PD-1-OX40, FasBB), and dual-CAR single-Switch (FasBB) configurations (single-switch FasBB vector pairs with CD19-CAR vector). **d-g**, Flow cytometry analysis of Zip-sorted BALB/c T cells transduced with dual-CAR and dual-CAR FasBB vectors or with multi-Switch vectors and stimulated for 24h with BM185-CD19 (E:T = 1:2) or left unstimulated. Data are mean ± SEM (biological replicates) of triplicate wells from n=4 donor experiments. Statistical comparisons were calculated using two-way ANOVA, with Tukey’s test. **h**, Flow cytometry analysis of BM185-ffluc-Thy1.1-Neo cell lines engineered to express FasL, PD-L1, or CD200. **i-k**. Sublethally irradiated BALB/c mice were injected with 1:1:1 mixture of BM185-CD19-FasL, BM185-CD20-PD-L1, and BM185-CD20-CD200 and treated with Zip-sorted dual-CAR BALB/c T cells, pre-cultured for 2 days with 1 μM dasatinib. **i**, leukemia BLI, and **j**, mouse survival, respectively, from one experiment. Survival differences were compared via log-rank test. **k**, Flow cytometry analysis of inhibitory ligand expression of single-ligand-positive BM185 lines harvested from bone marrow of mice reaching humane endpoints. PD-L1 expression difference was compared via a two-tailed t-test. **l**, Live-cell microscopy (Incucyte) analysis of EGFP-labelled dual-CAR and dual-CAR triple-Switch Zip-sorted BALB/c T cells co-cultured with iRFP713^+^ BM185 target cell lines as depicted (E:T = 1:1), with repetitive target addition on days 0, 1, and 2. *p≤0.05, **p≤0.01, ***p≤0.001, ****p≤0.0001, NS p>0.05.

**Extended Data Fig. 7:**
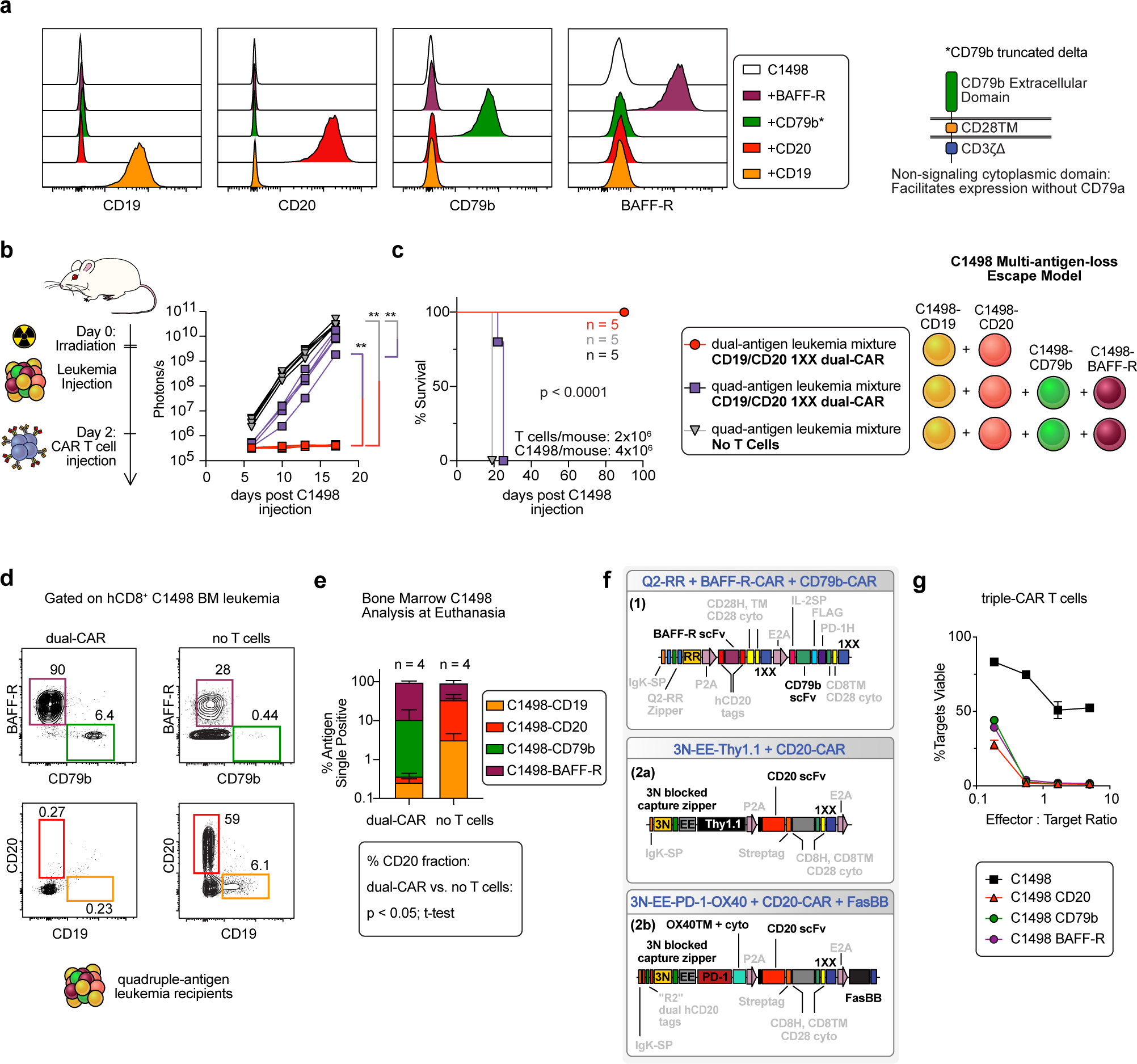
Dual-CAR T cells eliminate cognate targets and promote antigen-negative escape of leukemia mixture with high antigen heterogeneity. **a**, C1498 acute myeloid leukemia cell line was modified to singly express murine CD19, CD20, CD79bΔ (CD79b extracellular domain fused to CD28TM and CD3ζΔ; to promote surface expression without CD79a), and BAFF-R. **b-e**. Albino B6 mice were sublethally irradiated and injected with either 1:1 mixture of C1498-CD19 and C1498-CD20 or a 1:1:1:1 mixture of C1498-CD19, of C1498-CD20, C1498-CD79b, and C1498-BAFF-R and treated with dasatinib-cultured Zip-sorted dual-CAR 1XX T cells or left untreated. **b**, Leukemia BLI from single experiment. **c**, Survival. Log-transformed BLI AUC values were compared using a Vardi test with FDR correction. Survival differences were compared with a log-rank test. **d**, Target antigen expression of representative C1498 leukemia harvested from bone marrow at time of euthanasia for leukemia progression. Gated on hCD8^+^ C1498. **e**, Target antigen expression on bone marrow leukemia obtained from mice reaching humane endpoints. % CD20^+^ C1498 fraction of total bone marrow C1498 was compared with two-tailed t-test. **f**, Vector maps for constructs depicted in Fig. 5a. **g**, 24h luciferase-based target lysis assay with Zip-sorted BAFF-R/CD79b/CD20 triple-CAR BALB/c T cells with targets as depicted. Data are mean ±SEM of triplicate wells from single experiment. *p≤0.05, **p≤0.01, ***p≤0.001, ****p≤0.0001, NS p>0.05.

**Extended Data Fig. 8:**
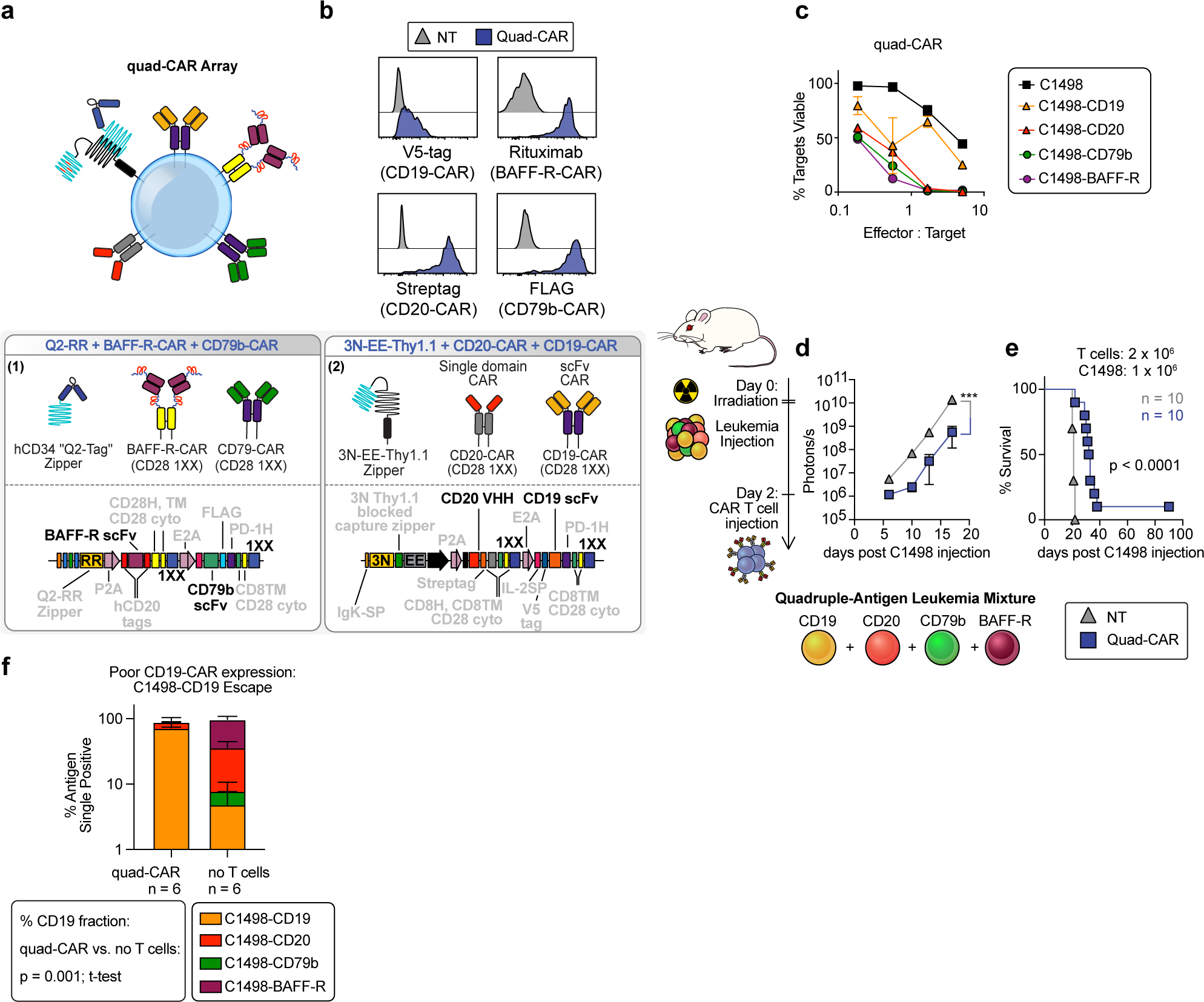
Poor expression of CD19-CAR in quad-CAR array promotes escape of CD19-positive C1498 leukemia. **a**, Diagram and vector maps for quad-CAR T cell receptor array. **b**, Flow cytometry analysis of CAR expression. **c**, 24h luciferase-based target lysis assay with Zip-sorted quad-CAR B6 T cells with targets as depicted. Data are mean ±SEM of triplicate wells from a representative experiment from n=2 donor replicates. **d-f**. Albino B6 mice were sublethally irradiated and injected with a 1:1:1:1 mixture of C1498-CD19, of C1498-CD20, C1498-CD79b, and C1498-BAFF-R and treated with dasatinib-cultured Zip-sorted quad-CAR 1XX T cells or left untreated. **d**, Leukemia BLI from two combined experiments. **e**, Survival. Log-transformed BLI AUC values were compared using a Vardi test with FDR correction. Survival differences were compared with a log-rank test. **f**, Target antigen expression of C1498 leukemia harvested from bone marrow at time of euthanasia for leukemia progression. Gated on hCD8^+^ C1498. % CD19^+^ C1498 fraction of total bone marrow C1498 was compared with two-tailed t-test. *p≤0.05, **p≤0.01, ***p≤0.001, ****p≤0.0001, NS p>0.05.

**Extended Data Fig. 9:**
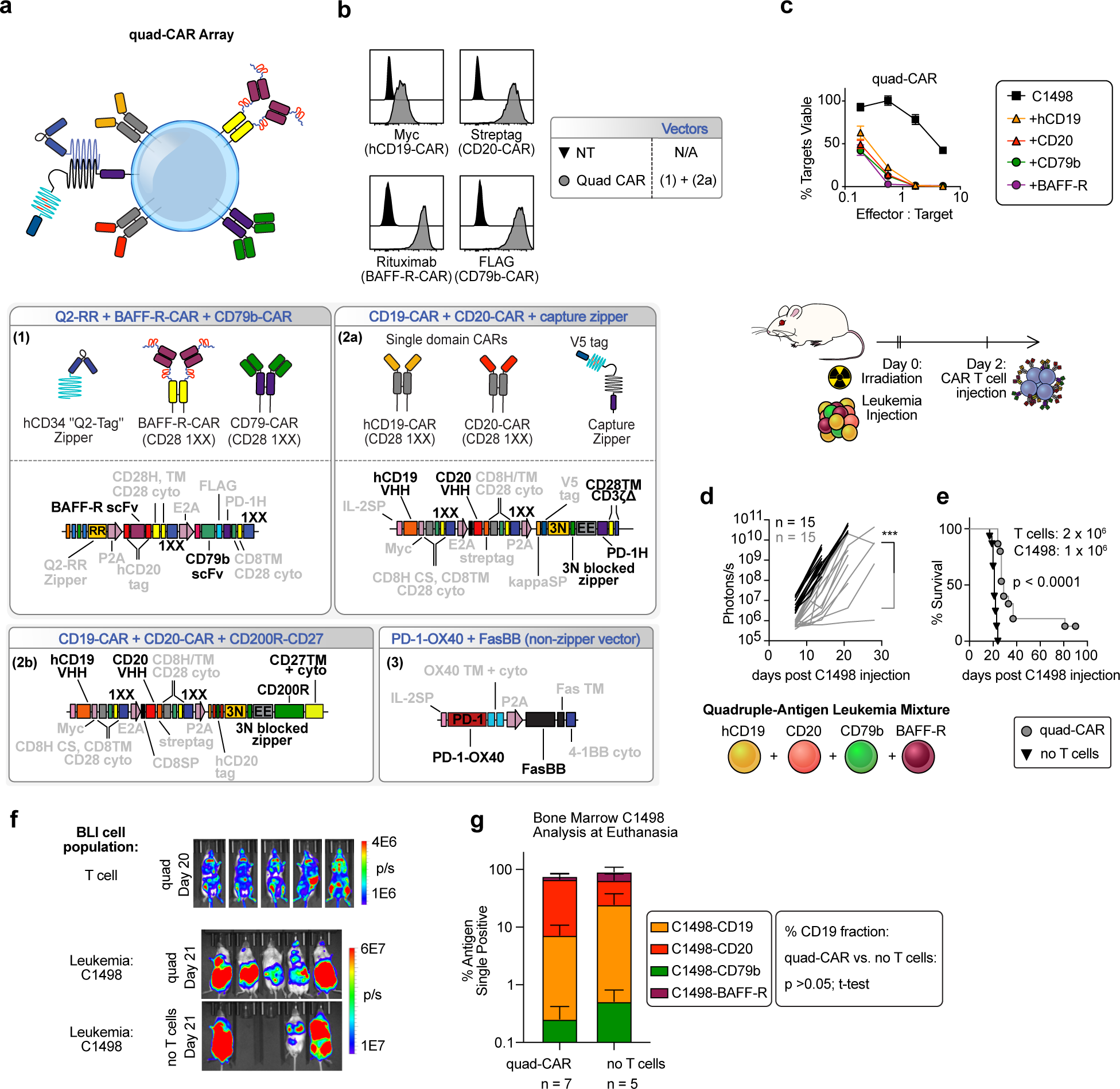
Quad-CAR T cells utilizing single-domain CD19-CAR demonstrate cognate target lysis, but exhibit poor anti-leukemia activity in vivo. **a**, Diagram and vector maps for quad-CAR receptor arrays. **b**, Flow cytometry analysis of CAR expression. **c**, 24h luciferase-based target lysis assay with quad-CAR B6 T cells and targets as listed. Data are mean ± SEM of triplicate wells for representative experiments from n=2 donors. **d-g**, Sublethally irradiated albino B6 mice were injected with 1:1:1:1 ratio of C1498 singly expressing (hCD19, CD20, CD79bΔ, BAFF-R) and treated with dasatinib-cultured quad-CAR albino B6 T cells co-transduced with Gaussia luciferase vector gLuc-PD-1H-CD24-GPI-P2A-EGFP. **d**, C1498 BLI from three combined experiments with different BLI imaging timing. **e**, Survival. BLI AUC values were compared using a Vardi test with FDR correction. Survival differences were compared via pairwise log-rank test. **f**, Representative day 20 T cell BLI (Gaussia) and day 21 C1498 BLI (CBR). **g**, Antigen expression analysis from C1498 harvested from bone marrow of mice reaching humane endpoints. % hCD19^+^ C1498 was compared with two-tailed t-test. *p≤0.05, **p≤0.01, ***p≤0.001, ****p≤0.0001, NS p>0.05.

**Extended Data Fig. 10:**
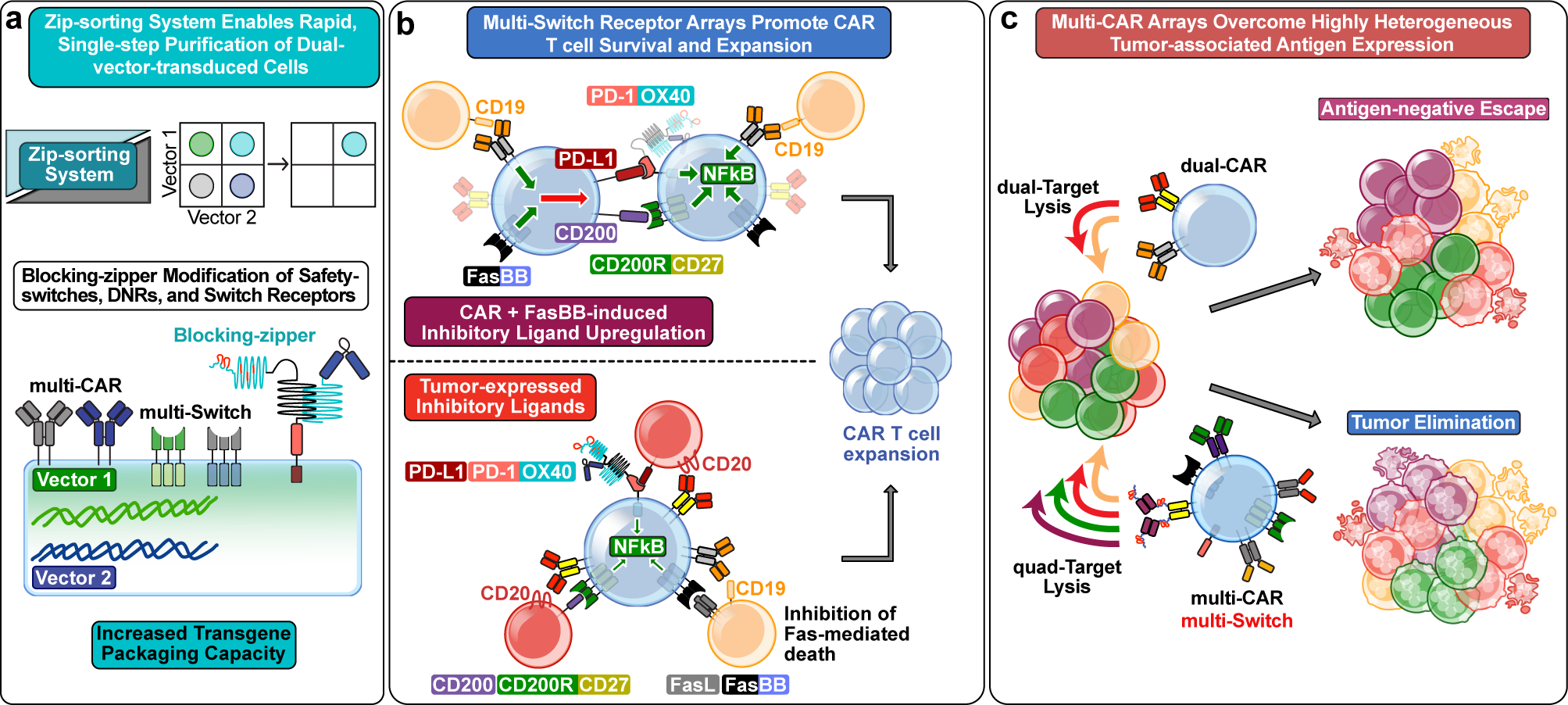
Summary of Zip-sorting system and multi-CAR multi-Switch T cells. **a**, Zip-sorting system directs single-step immunomagnetic purification of dual-vector-transduced cells. The blocking-zipper strategy enables zipper modification of safety-switches, DNRs, switch receptors, and CARs by promoting preferential surface expression of the affinity-tag zipper on dual-transduced cells. Supporting data: Fig. 1 and Extended Data Fig. 1. **b**, (Top) Proposed mechanism for enhanced proliferation of multi-Switch CAR T cells in response to leukemia targets without inhibitory ligand overexpression. Antigen stimulation drives upregulation of PD-L1 and CD200 on dual-CAR FasBB switch receptor-expressing T cells. PD-1-OX40 and CD200R-CD27 switch receptors signal in response to T cell-expressed inhibitory ligands. Supporting data: Fig. 4e and Extended Data Fig. 6a-b, d-g. (Bottom) Proposed mechanism for enhanced proliferation and survival of multi-Switch CAR T cells in response to leukemia targets with inhibitory ligand overexpression. Direct stimulation of switch receptors via tumor cell-expressed inhibitory ligands promotes TNFRSF signaling. FasBB inhibits Fas/FasL-mediated T cell death (dominant negative function). Supporting data: Extended Data Fig. 6l. **c**, Dual-CAR T cells promote immunoediting and antigen-negative outgrowth of leukemia in response to a leukemia population heterogeneously expressing four tumor-associated antigens. Supporting data: Extended Data Fig. 7. Triple-CAR and quad-CAR T cells target greater number of antigens and can overcome higher antigen heterogeneity. Co-expression of multiple switch receptors promotes proliferation and enhances anti-leukemia activity of multi-CAR T cells. Supporting data: Figs. 5-6.

